# FOXM1 expression reverts aging chromatin profiles through repression of the senescence-associated pioneer factor AP-1

**DOI:** 10.1101/2023.05.04.539315

**Authors:** Fábio J. Ferreira, Mafalda Galhardo, Joana Teixeira, Elsa Logarinho, José Bessa

## Abstract

Aging is characterized by changes in gene expression that drive deleterious cellular phenotypes leading to senescence. The transcriptional activation of senescence genes has been mainly attributed to epigenetic shifts, but the changes in chromatin accessibility and its underling mechanisms remain largely elusive in natural aging. Here, we profiled chromatin accessibility in human dermal fibroblasts (HDFs) from neonatal and octogenarian individuals. We found that AP-1 binding motifs are prevalent in elderly specific accessible regions of the chromatin while neonatal-specific regions are highly enriched for TEAD binding motifs. We further show that *TEAD4* and *FOXM1* share a conserved transcriptional regulatory landscape controlled by an age-dependent enhancer that closes with aging and drives senescence when deleted. Finally, we demonstrate that *FOXM1* ectopic expression in elderly cells partially resets chromatin accessibility to a youthful state due to FOXM1 repressive function in the promoters of several members of the AP-1 complex. These results place *FOXM1* at a top hierarchical level in chromatin remodeling required to prevent senescence.

## INTRODUCTION

Aging is characterized by the time-dependent functional decline that affects virtually all organisms. Transcriptional signatures of natural aging and lifespan extension have been identified (Stegeman & Weake, 2017; Tyshkovskiy *et al*, 2019). Epigenetic changes have been explored as part of the mechanisms controlling the age-related transcriptional alterations, either by measuring the total load of histone modifications in human aging cells (Cheung *et al*, 2018) or by examining the genome-wide distribution of specific histone marks associated to transcriptional regulatory functions (Zhou *et al*, 2018). Moreover, chromatin accessibility profiling in cellular models of induced senescence that partly recapitulate natural cellular aging has defined the dynamics and organizational principles of cis-regulatory elements (CREs) driving senescence transcriptional programs (Martínez-Zamudio *et al*, 2020; Zhang *et al*, 2021). One key player that imprints the senescence cis-regulatory signature is the activator protein 1 (AP-1) complex (Martínez-Zamudio *et al*, 2020; Zhang *et al*, 2021; Di Giorgio *et al*, 2021; Wang *et al*, 2022). Importantly, impairment of the pioneer transcription factor (TF) JUN, a member of the AP-1 complex, partially reverts a transcriptional programme of senescent cells (Martínez-Zamudio *et al*, 2020). These results suggest that increased AP-1 function leads to a senescent cellular state, although the mechanisms controlling AP-1 expression remain unknown. In the opposite spectrum of senescence regulating factors, YAP:TEAD and FOXM1 transcription factors have been shown to play key roles in the maintenance of cellular fitness and proliferation. YAP is normally sequestered in the cytoplasm by Hippo pathway growth-inhibitory signals, but in response to mitogenic signaling, undergoes nuclear translocation and binding to the TEAD1-4 family of TFs (Boopathy & Hong, 2019). YAP:TEAD physically interacts with B-MYB:MuvB (MMB) and FOXM1 to promote the expression of G2/M cell cycle genes through long-range interactions on chromatin between YAP:TEAD at enhancers and MMB:FOXM1 at promoters (Pattschull *et al*, 2019; Gründl *et al*, 2020). Both *YAP* and *FOXM1* downregulation drive transcriptional signatures of aging (Xie *et al*, 2013; Macedo *et al*, 2018), and, conversely, *YAP* and *FOXM1* sustained expressions are able to delay senescence and its associated traits in animal models of physiological and accelerated aging (Fu *et al*, 2019; Ribeiro *et al*, 2022; Sladitschek-Martens *et al*, 2022). However, the mechanisms behind *YAP* and *FOXM1* downregulation during aging, and the crosstalk with the AP-1-mediated cis-regulatory signature driving an aging-associated senescence transcriptome, are yet to be identified.

In this work, we profiled chromatin accessibility by performing Assay for Transposase-Accessible Chromatin using sequencing (ATAC-seq) (Buenrostro *et al*, 2013) in early passage human dermal fibroblasts (HDFs) retrieved from neonatal and octogenarian healthy individuals. We found that elderly-specific accessible regions of the chromatin are enriched for AP-1 binding motifs, while neonatal-specific regions are enriched for TEAD motifs, which bind TFs of the Hippo signaling pathway. Interestingly, we found that changes in chromatin accessibility during aging correlate with transcriptional changes in regulatory landscapes. One example is the transcriptional regulatory landscape comprising *TEAD4* and *FOXM1*, which we found to modulated by an enhancer whose accessibility is lost with aging. Excitingly, we demonstrated that FOXM1 repressive function in the promoters of several members of the AP-1 complex is required to sustain chromatin accessibility at a youthful state. Thus, we bring mechanistic insight into FOXM1 repression during natural aging and disclose its top hierarchical function in rescuing AP-1-driven senescence.

## RESULTS

### Changes in chromatin accessibility during cellular aging are associated with enrichment of AP-1-binding motifs and loss of TEAD-binding motifs

To understand the impact of natural aging in the accessibility of chromatin profiles in a genome-wide manner, we have performed ATAC-seq in neonatal and elderly HDFs at early cell culture passage, with population doublings well below replicative exhaustion (Macedo *et al*, 2018). We found 20.491 sequences that are accessible specifically in neonatal cells (neonatal-specific;), 32.012 sequences accessible specifically in elderly cells (elderly-specific) and 62.836 sequences shared by neonatal and elderly cells (common; Figure 1a-c). We have further analyzed the ATAC signal in the group of common sequences. Out of the 62.836 common sequences, 2.083 were enriched in neonatal cells and 4.779 were enriched in elderly cells (Log2FC>1; adjusted *p* value<0.05; Figure 1c). We intersected the ATAC-seq peaks with available ENCODE candidate cis-regulatory elements (cCREs) (Moore *et al*, 2020) (Supplementary Table 1) and found that about 60% of peaks from both ages overlap with distal enhancer-like elements. This percentage is maintained among age-specific and common elements, but the overlap is bigger among common sequences with age-specific accessibility enrichment (67% in neonatal-enriched; 83% in elderly-enriched). In contrast, we found that the percentage of overlap with promoter-like and proximal enhancer-like elements is smaller in age-specific sequences than in common sequences (20% of common, 7% of neonatal-specific and 3% of elderly-specific sequences overlap with promoter-like elements; 21% of common, 13% of neonatal-specific and 11% of elderly-specific sequences overlap with proximal enhancer-like elements; Supplementary Table 1). These data suggest that most age-associated changes in chromatin accessibility in HDFs occur in distal enhancer-like elements, that typically regulate multiple genes. Next, we performed a motif enrichment search for TF binding sites in neonatal and elderly specific open chromatin regions. We analyzed and compared the enrichment of each TF motif in elderly and neonatal-specific regions (Figure 1d; Supplementary Table 2). We observed that binding motifs for *TEAD* proteins were less enriched in elderly-specific sequences (Δ enrichment: *TEAD3:* -10,66%; *TEAD1*: -10,23%; *TEAD4: -8,48%* and *TEAD2: -6,23%*), while members of the AP-1 complex were more enriched in elderly specific sequences (Δ enrichment: *JUN*: 24,20%; *ATF3*: 24%; *FOS: 22,90%*; and *JUNB: 21,63%;* Figure 1d). Similar results were obtained when taking into account only the subset of neonatal and elderly-specific open chromatin regions that presented significant quantitative differences of ATAC-seq signal or when considering all age specific and age-enriched regions (Log2FC>1; adjusted *p*-value <0.05; Supplementary Figure 1a,b; Supplementary Table 3 and 4). These results suggest that TEAD and AP-1 TFs exhibit inversely correlated dynamics during aging and are in agreement with previous studies reporting the enrichment of AP-1 binding motifs in open chromatin regions of induced senescent cells (Martínez-Zamudio *et al*, 2020; Zhang *et al*, 2021), as well as TEAD-mediated senescence inhibition by the Hippo pathway (Xie *et al*, 2013).

**Figure 1.**
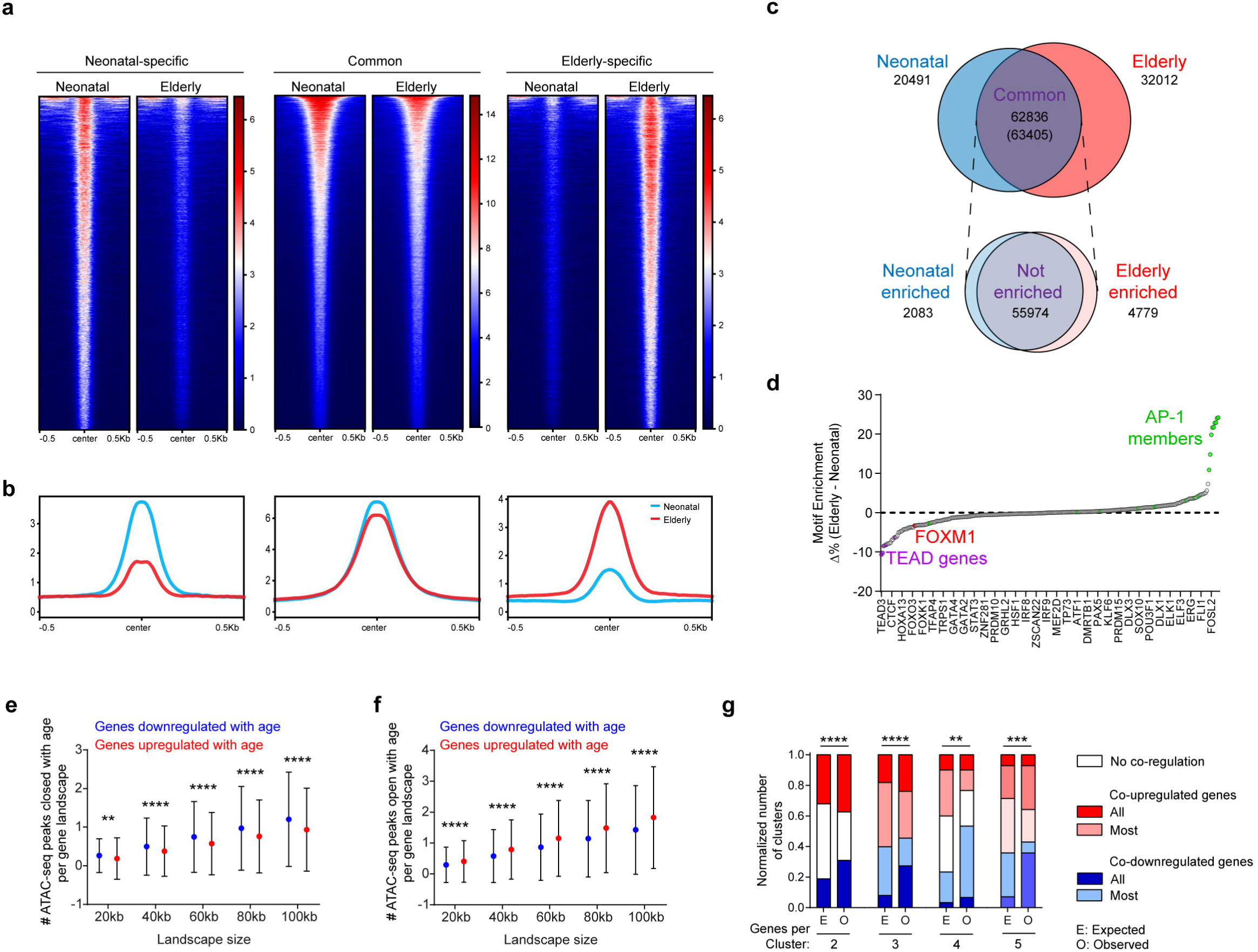
Chromatin accessibility variation during aging. a. Heatmaps showing ATAC-seq signals across age-specific and common reproducible peaks in neonatal and octogenarian HDFs. Each peak was ordered in descending order based on the mean value of ATAC-seq signal per region. b. Profile plots for scores over sets of age-specific and common ATAC-seq reproducible peaks in neonatal and octogenarian HDFs. The Y axis represents mean aligned reads per-base from two replicates. c. Venn diagram showing overlaps of the neonatal and elderly-specific ATAC-seq reproducible peaks. *Top:* All peaks defined with MACS2 and selected with IDR. Common peaks refer to neonatal peaks with overlap with at least one elderly peak. Value between parenthesis represents the elderly peaks with overlap with at least one neonatal peak. *Bottom:* Common peaks with significantly enriched signal in either neonatal or elderly samples, as determined by DESeq2 analysis. d. Comparison of HOMER-defined Known Motif Enrichment between elderly and neonatal-specific ATAC-seq reproducible peaks. Negative values represent enrichment in neonatal peaks, while positive values represent enrichment in elderly peaks. Horizontal axis names only every other 9 TFs, for simplicity. e,f. Quantification of ATAC-seq reproducible peaks whose accessibility changes between neonatal and elderly cells, in genes that are up (red) or downregulated (blue) during aging. e shows peaks that close during aging (neonatal-specific). f shows peaks that open during aging (elderly-specific). Values represent mean ± sd. *p*-value was determined by Mann-Whitney U test. g. Quantification of observed gene clusters, 2 to 5 genes per cluster, showing co regulation (either up or downregulation) or no co-regulation of gene expression during aging. *p*-value was determined by Chi-square test between the expected (E) and observed (O) number of clusters per class.

### Changes in chromatin accessibility during aging correlate with transcriptional shifts in regulatory landscapes

To evaluate the relevance of the chromatin state during aging, we correlated the accessibility of age-specific regions with the transcriptional output of nearby genes. We analyzed available transcriptomic datasets for HDFs retrieved from the same neonatal and elderly donors, cultured under the same conditions (Macedo *et al*, 2018) and we established two sets of genes, either up or downregulated during aging. For each gene of the up or downregulated set, we quantified the number of neonatal-specific (closed with age) and elderly-specific (open with age) ATAC-seq peaks that are located in their genomic vicinity (from 20kb to 100kb). We observed that the number of neonatal-specific peaks is higher in the genomic landscapes of downregulated genes than in upregulated genes with aging (Figure 1e). Conversely, the number of elderly-specific peaks is higher in the genomic landscapes of upregulated genes than in downregulated genes with aging (Figure 1f). Importantly, in both comparisons, these differences became more striking when analyzing larger genomic landscapes, up to 100kb centered in each gene (Figure 1e,f). It is noteworthy that the number of age-independent accessible peaks between genes that are up and downregulated during aging was not significantly different (Supplementary Figure 1c). Altogether, these results suggest that genes whose expression changes with advancing age are under the control of CREs, which in turn are modulated by changes in chromatin accessibility during aging. Moreover, the difference in the average number of ATAC-seq peaks nearby up and downregulated genes increases with the extended span of the genomic landscapes (from 20kb to 100kb), both for neonatal and elderly-specific peaks, pointing to the existence of large genomic landscapes where multiple genes are transcriptionally coordinated by common CREs during aging. To explore this hypothesis, we defined clusters of genes that are no more than 100kb part from each other and we asked if they presented a similar transcriptional change, either upregulation or downregulation during aging, in comparison to a theoretical assumption of no co-regulation. We observed that the gene clusters, regardless of their size (2 to 5 genes), present coordinated transcriptional changes during aging (Figure 1g; Supplementary Table 5), further supporting the existence of age-associated regulatory landscapes.

### *TEAD4*, *FOXM1* and *RHNO1* belong to the same regulatory landscape

To understand if the coordinated transcriptional changes of several genes within a regulatory landscape could be determinant for aging, we explored the list of putative aging-associated regulatory landscapes (Supplementary Table 5). We focused our attention on the landscape containing *TEAD4*, a member of the *TEAD* gene family that encodes TFs whose binding motifs were the most enriched in neonatal-specific ATAC seq peaks (Δ enrichment˃10%; Figure 1d; Supplementary Table 2). TEAD4 is known to regulate cell proliferation, tissue growth and apoptosis (Huang *et al*, 2005; Watt *et al*, 2017) and its repression has been associated with senescence in human cells (Fu *et al*, 2019). Within the same regulatory landscape, we found *FOXM1*, a gene encoding a TF required for a wide spectrum of essential biological functions, including DNA damage repair and cell proliferation (Laoukili *et al*, 2007; Zona *et al*, 2014), and whose ectopic expression has been shown to revert senescence phenotypes in *in vitro* and *in vivo* models of premature and natural aging (Macedo *et al*, 2018; Ribeiro *et al*, 2022). Importantly, the FOXM1 binding motif was highly represented in neonatal-specific ATAC seq peaks (Δ enrichment=3.28%; Figure 1d; Supplementary Table 2). Furthermore, in the close genomic vicinity of *FOXM1* we found *RHNO1*, which has been implicated in DNA replication stress response (Cotta-Ramusino *et al*, 2011; Hara *et al*, 2020). *FOXM1* and *RHNO1* exist in a head-to-head orientation, sharing a bidirectional promoter (Barger *et al*, 2021), further supporting that the genes within this landscape might share regulatory information. Moreover, except for *ITFG2*, all the genes within this putative regulatory landscape (*FKBP4*, *FOXM1*, *RHNO1*, *TULP3*, *TEAD4* and *TSPAN9*; no information for *NRIP2* and *TEX52*) were found downregulated in elderly cells (Supplementary Figure 2a), suggesting that the co-expression of these genes might sustain a youthful cell fitness. Promoter DNA methylation (Rakyan *et al*, 2010; King *et al*, 2012; Ehrlich, 2019) was ruled out as the causal mechanism behind the aging associated coordinated transcriptional downregulation of the *TEAD4/FOXM1/RHNO1* gene cluster since we found the *FOXM1/RHNO1* promoter to be hypomethylated in both neonatal and elderly HDFs (*p*-value=0.964; Supplementary Figure 2b). To assess co regulation by shared CREs as an alternative mechanism, we pursued with a 4C-seq analysis of chromatin interactions within this gene cluster in neonatal HDFs. We selected the bidirectional promoter of *FOXM1/RHNO1*, roughly located in the middle of the gene cluster, as the viewpoint for the 4C-seq experiment. We found interactions of the *FOXM1/RHNO1* promoter with several genomic regions, including promoters of nearby genes, spanning approximately 300kb (Supplementary Figure 2c), thereby supporting transcriptional co-regulation of genes within this regulatory landscape by shared distal CREs.

Interestingly, and attesting its functional relevance, the *TEAD4/FOXM1/RHNO1* gene cluster is conserved in several vertebrate lineages, including distant Gnathostomata species (about 473 million years) (Supplementary Figure 3a) (Nguyen *et al*, 2022; Bateman *et al*, 2023). Non-vertebrate species such as lancelets also present the bidirectional promoter configuration of *FOXM1* and *RHNO1* (or *RHNO1*-like) (Yu *et al*, 2008; Aldea *et al*, 2015; Marlétaz *et al*, 2018; Nguyen *et al*, 2022) (Supplementary Figure 3b,c). Moreover, the tunicate *Ciona intestinalis* presents a bidirectional promoter co-regulating *FOXM1* and a CCNB1IP1-like gene (Cunningham *et al*, 2022; Bateman *et al*, 2023), whose E3 ubiquitin-protein ligase function is shared with the human 9-1-1 complex comprising RHNO1 (Cotta-Ramusino *et al*, 2011; Lindsey-Boltz *et al*, 2015) (Supplementary Figure 3d). These data suggest this genomic organization dates back about 550 million years and it is functionally advantageous to all Chordata (Nguyen *et al*, 2022) (Supplementary Figure 3e).

### C10 is an age-dependent CRE that controls the expression of *TEAD4*, *FOXM1* and *RHNO1*

Using the ATAC-seq datasets described above, we found many accessible chromatin regions in the genomic landscape of *FOXM1* and *RHNO1* (Figure 2a). The promoter regions of *FOXM1*/*RHNO1* and of the nearby genes overlap with open chromatin regions, supporting active gene transcription in neonatal HDFs (Macedo *et al*, 2018). Many accessible loci are localized in inter and intragenic regions, pointing to the existence of functional non-coding CREs regulating gene expression within the landscape. We then crossed our datasets from neonatal HDFs (4C-seq and ATAC-seq) with ChIP-seq data for H3K27ac, H3K4me3 and transcription factor clusters (TF clusters) from multiple cell types from the ENCODE project (Rosenbloom *et al*, 2013; Navarro Gonzalez *et al*, 2021) to explore putative functions of the open chromatin regions interacting with the *FOXM1*/*RHNO1* promoter. This combined analysis allowed us to select 14 putative CREs, which we designated C1 through C14 (Figure 2a; Supplementary Table 6). All selected putative CREs include at least one open chromatin region (ATAC-seq) and very low levels of the H3K4me3 mark, thus excluding likely promoter regions (Liang *et al*, 2004). We then performed luciferase reporter assays to test the enhancer activity of the 14 putative CREs in neonatal HDFs (Figure 2b). We found regions C2, C5, C9 and C10 to induce significant luciferase activity in comparison to control, thus indicating that the *FOXM1*/*RHNO1* promoter interacts with active enhancers in neonatal HDFs. To determine if these enhancers contribute to *FOXM1* expression in these cells, we performed CRISPR/Cas9-mediated genomic deletions (Figure 2c) of the regions C2, C5, C9 and C10, as well as of the region C8, the closest open chromatin region to the *FOXM1*/*RHNO1* promoter without enhancer activity. Deletion of enhancers C5, C9 and C10 significantly downregulated *FOXM1* expression in polyclonal populations of neonatal HDFs retrieved by FACS (Figure 2d). As expected, deletion of the C8 region had no impact on gene expression. These results suggest that enhancers C5, C9 and C10 are required to sustain *FOXM1* expression in neonatal HDFs. To examine if changes in the interactome of *FOXM1*/*RHNO1* promoter account for the *FOXM1* downregulation previously observed along aging (Macedo *et al*, 2018), we then performed 4C-seq in elderly HDFs (Supplementary Figure 4a). We found the interactome of the *FOXM1*/*RHNO1* promoter to be kept similar in neonatal and elderly HDFs. Thus, conformational changes at the *FOXM1*/*RHNO1* regulatory landscape unlikely explain the transcriptional shift observed in elderly cells. In contrast, we found many ATAC-seq peaks to be specific to either neonatal or elderly cells, suggesting that alterations in accessible chromatin may contribute to the age-associated transcriptional changes (Supplementary Figure 4b). Hence, we took a closer look on the ATAC-seq peaks of the enhancer regions C2, C5, C9 (Supplementary Figure 4c) and C10 (Figure 3a) to explore differences between neonatal and elderly HDFs. We found chromatin accessibility to be retained in enhancers C2, C5, C9, but not in enhancer C10, in elderly HDFs. Enhancer C10 comprises two open chromatin regions in neonatal HDFs, one of which becomes closed in elderly HDFs (Figure 3a). We designated the age-independent region as C10 P1 and the age-dependent region as C10-P2. The deletion of either C10-P1 or C10-P2 in neonatal HDFs resulted in significant downregulation of *FOXM1* and *RHNO1* expression (Figure 3b.c), suggesting that both C10 regions are functionally required for proper gene expression. We also assessed the impact on *TEAD4* expression, observing that the deletion of either C10-P1 or C10-P2 induces significant downregulation of *TEAD4* (Figure 3d). These results demonstrate that the age-dependent C10 enhancer coordinates the expression of *TEAD4*, *FOXM1* and *RHNO1*.

**Figure 2.**
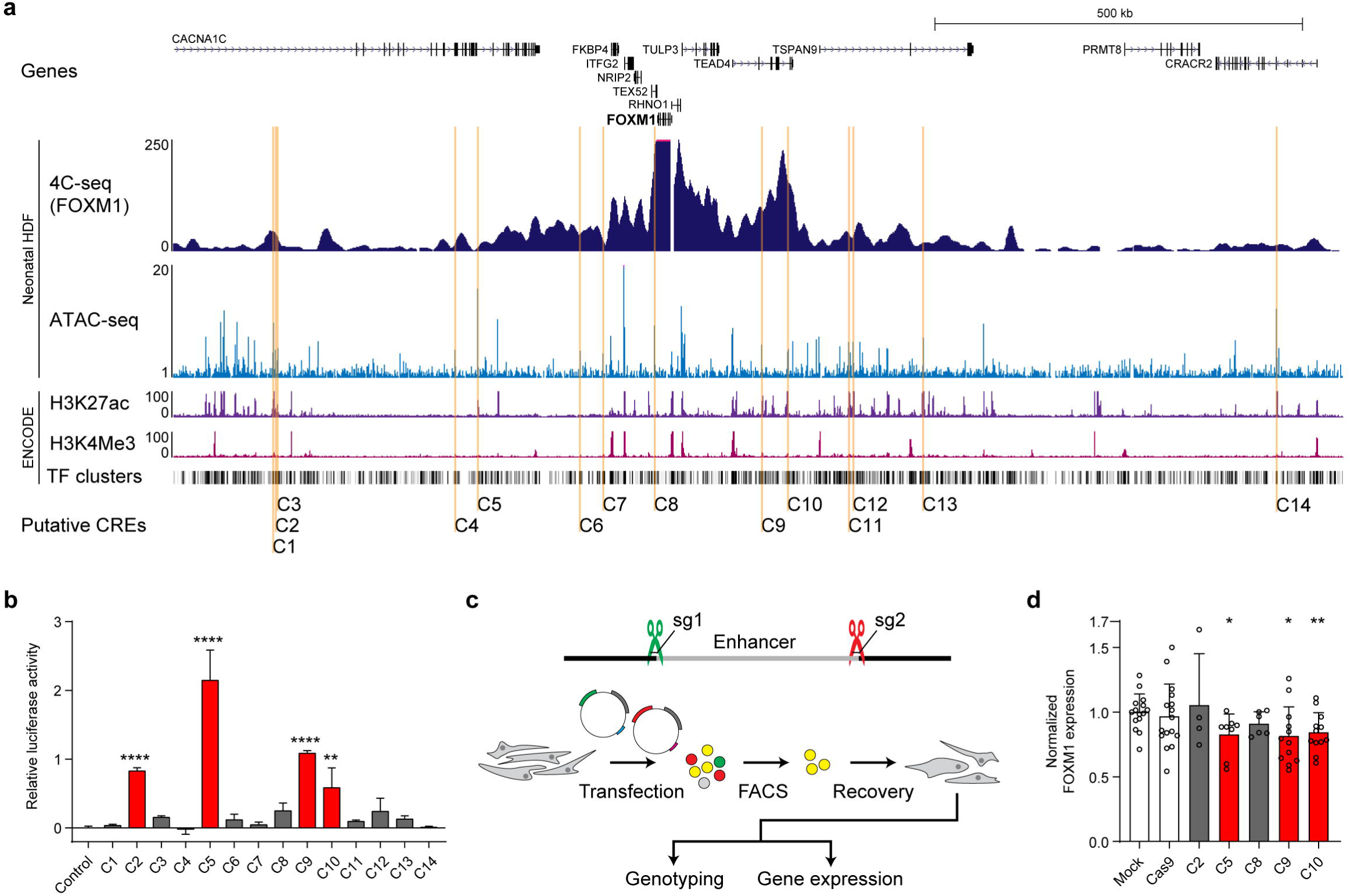
Non-coding genomic regions with enhancer activity are required for normal *FOXM1* expression. a.Genomic tracks illustrating epigenetic features nearby *FOXM1/RHNO1*. 4C-seq data (dark blue) and ATAC-seq data (light blue) reveal non-coding genomic regions interacting with the *FOXM1/RHNO1* promoter that are also accessible, thus likely active, in neonatal HDFs. ChIP-seq data for H3K27ac (purple) and H3K4Me3 (pink), and TF clusters, all from the ENCODE project, allowed the selection of putative cis-regulatory elements. Based on these attributes, 14 putative cis-regulatory elements (orange) were selected for further examination. b.Luciferase reporter assay for the 14 putative enhancer regions in neonatal HDFs. Red bars highlight sequences with significant enhancer activity. ** *p*-value ≤0.01; **** *p*-value ≤0.0001 by one-way ANOVA with Dunnett’s correction for multiple comparisons. Values represent mean ± SD. c.Experimental layout of CRISPR/Cas9-mediated genomic deletions of putative *FOXM1* enhancers. Neonatal HDFs were transfected with two vectors containing the Cas9 coding sequence, each carrying a different fluorescent protein coding gene and a sgRNA targeting either the upstream or downstream edge of the enhancer sequence. Double positive cells were FACS-sorted, plated, and allowed to recover. Genomic DNA for genotyping and RNA for gene expression analysis were collected simultaneously. d.Gene expression analysis of *FOXM1* by RT-qPCR upon CRISPR/Cas9-mediated deletion of enhancers C2, C5, C9 and C10 and control region C8 in neonatal HDFs. Mock transfection and transfection with Cas9 alone (Cas9) were used as controls. Deletion of C5, C9 or C10 lead to significant FOXM1 downregulation (red bars). * *p*-value ≤0.05; ** *p*-value ≤0.01 by unpaired Student’s t-test. Values represent mean ± SD.

**Figure 3.**
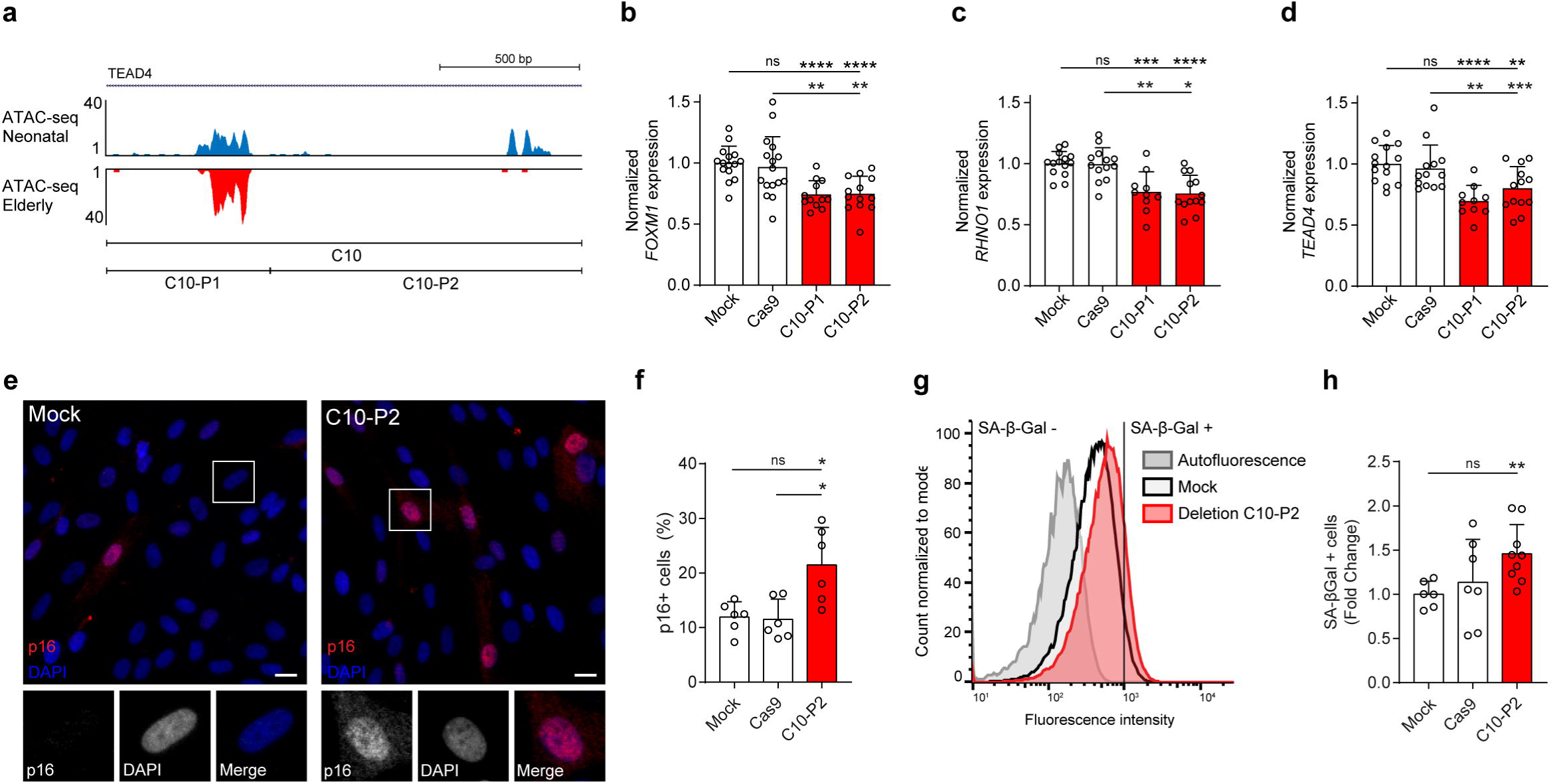
Deletion of enhancer with age-specific chromatin accessibility leads to gene expression changes and senescence phenotypes. a. ATAC-seq profile of enhancer C10, located in a *TEAD4* intron, in neonatal (blue) and octogenarian (red) HDFs. One region (C10-P1) is kept accessible while other (C10-P2) is closed in elderly cells. b-d. Gene expression analysis of *FOXM1* (b), *RHNO1* (c) and *TEAD4* (d) by RT-qPCR in neonatal HDFs carrying genomic deletion of C10-P1 or C10-P2 (red) compared to mock control cells (white) and Cas9-only treated cells (white). * *p*-value ≤0.05; ** *p*-value ≤0.01; *** *p*-value ≤0.001; **** *p*-value ≤0.0001 by unpaired Student’s t-test. Values represent mean ± SD. c.p16/CDKN1A immunostaining (red) (and DNA counterstaining with DAPI, blue) in mock control neonatal HDFs (left) and neonatal HDFs with deletion of C10-P2 (right). Insets highlights a p16-negative control cell (left) and a p16-positive cell (right). Scale bar: 20 μm. d.Quantification of the percentage of C10-P2-deleted cells staining positive for the p16 senescence marker (red) compared to mock control cells (white) and Cas9-only treated cells (white). * *p*-value ≤0.05 by unpaired Student’s t-test. Values represent mean ± SD. e.Cytometry plot representing the frequency and intensity of SA-β-Gal activity in mock treated cells (black outline) vs. C10-P2-deleted cells (red outline). A vertical line divides the intensities considered negative (SA-β-Gal-) and positive (SA-β-Gal+) for SA-β-Gal activity, based on the autofluorescence control (grey outline). f.Fold-change in the number of cells positive for SA-β-Gal activity between mock control cells, Cas9-only treated cells and cells lacking C10-P2 (red). ** *p*-value ≤0.01 by unpaired Student’s t-test. Values represent mean ± SD.

### C10 deletion is sufficient to trigger senescence in neonatal HDFs

Since C10-P2 accessibility is lost in naturally aged cells, we asked if the deletion of this region in neonatal HDFs is sufficient to trigger senescence, through *TEAD4*/*FOXM1*/*RHNO1* gene cluster downregulation. As previously reported for *TEAD4, FOXM1*, and *RHNO1* repressions (Laoukili et al, 2005; Kim et al, 2010; Cotta Ramusino et al, 2011; Zona et al, 2014; Lindsey-Boltz et al, 2015; Takeuchi et al, 2017; Macedo et al, 2018; Hazan et al, 2019), we found that CRISPR/Cas9-mediated deletion of C10-P2 induced a decreased percentage of cells staining positive for the Ki67 proliferation marker (Sobecki *et al*, 2017) (Supplementary Figure 5a,b), and an increased percentage of cells staining positive for 53BP1/p21, a combination of markers indicative of cell cycle arrest associated with DNA damage (Supplementary Figure 5c,d). While p21 activation might be only transient (Stein *et al*, 1999), expression of p16 accumulates with and is partly responsible for maintaining the cell cycle arrest associated with cell senescence (Alcorta *et al*, 1996). Further supporting that C10-P2 deletion drives senescence, we found an increased percentage of p16-positive cells (Figure 3e,f) and a higher activity of the lysosomal enzyme senescence-associated β-galactosidase (SA-β-gal) (Lee *et al*, 2006) (Figure 3f,h) in C10-P2-deleted vs. control cell populations. We additionally measured mitotic duration, which increases with aging due to *FOXM1* repression (Macedo *et al*, 2018), finding that C10-P2 deletion leads to mitotic delay (Supplementary Figure 5e). Overall, our data demonstrate that CRISPR/Cas9-mediated deletion of C10-P2 induces a set of senescence-associated phenotypes in neonatal HDFs. Altogether, these results suggest that the age-associated functional decline of an enhancer leads to a coordinated downregulation of (at least) *TEAD4, FOXM1* and *RHNO1,* three genes whose knockdown has been associated to senescence phenotypes.

### *FOXM1* overexpression in elderly HDFs rejuvenates the chromatin accessibility profile

Among the co-regulated senescence-associated triad of genes *TEAD4, FOXM1* and *RHNO1 FOXM1* stands out for its well-established role as modulator of aging phenotypes (Macedo *et al*, 2018; Ribeiro *et al*, 2022). We asked if this could be ascribed to a potential FOXM1 function in chromatin remodeling. To test this, we ectopically expressed a constitutively active truncated form of *FOXM1* (*FOXM1-dNdKEN*) (Laoukili *et al*, 2008b, 2008a; Macedo *et al*, 2018) in elderly HDFs (Supplementary Figure 6a,b) and performed ATAC-seq for chromatin profiling. We found that, after *FOXM1* ectopic expression, 25.016 peaks became inaccessible (26,2% of all elderly peaks) and 14.899 new accessible peaks were identified (17,5% of all peaks in elderly HDFs expressing FOXM1-dNdKEN; Supplementary Figure 6c). We then compared the peaks whose accessibility was altered by FOXM1 overexpression with the age-specific accessible peaks described above (neonatal-specific and elderly-specific; Figure 1c). We observed that 17.752 out of the 32.012 elderly-specific regions were closed (55,4%) and that, out of the 20.491 neonatal-specific regions, 6.488 were opened in elderly HDFs upon FOXM1 overexpression (31,7%; Figure 4a and Supplementary Figure 6d). Conversely, 88,9% of the regions whose availability is kept during aging (Common; Figure 1c) remains available upon *FOXM1* overexpression (55.849 out of 62.836; Supplementary Figure 6d). These results suggest that the ectopic expression of *FOXM1* partially shifts the chromatin accessibility profile towards a rejuvenated state. This shift is more evident in the number of closed chromatin regions (26,2% of all elderly regions, 55,4% of elderly specific regions) than in the number of opened regions (17,5% of all regions in FOXM1 overexpressing cells, 31,7% of neonatal-specific regions) in elderly HDFs as result of FOXM1 overexpression, suggesting that FOXM1 might predominantly repress factors that contribute to the establishment and maintenance of elderly-specific open chromatin regions. After performing motif discovery in elderly-specific regions that shift to an inaccessible state (closed) and in neonatal-specific regions that shift to an accessible state (opened) (Supplementary Figure 6d) and comparing the motif enrichment in each dataset, we found that components of the AP-1 complex are prevalent in the former whereas the CTCF motif is enriched in the latter (Figure 4b; Supplementary Table 7). Since AP-1 is known to function as a pioneer factor on the remodeling of senescence associated chromatin profiles (Martínez-Zamudio *et al*, 2020; Zhang *et al*, 2021; Di Giorgio *et al*, 2021; Wang *et al*, 2022), the results suggest that FOXM1 might downregulate members of the AP-1 complex.

**Figure 4.**
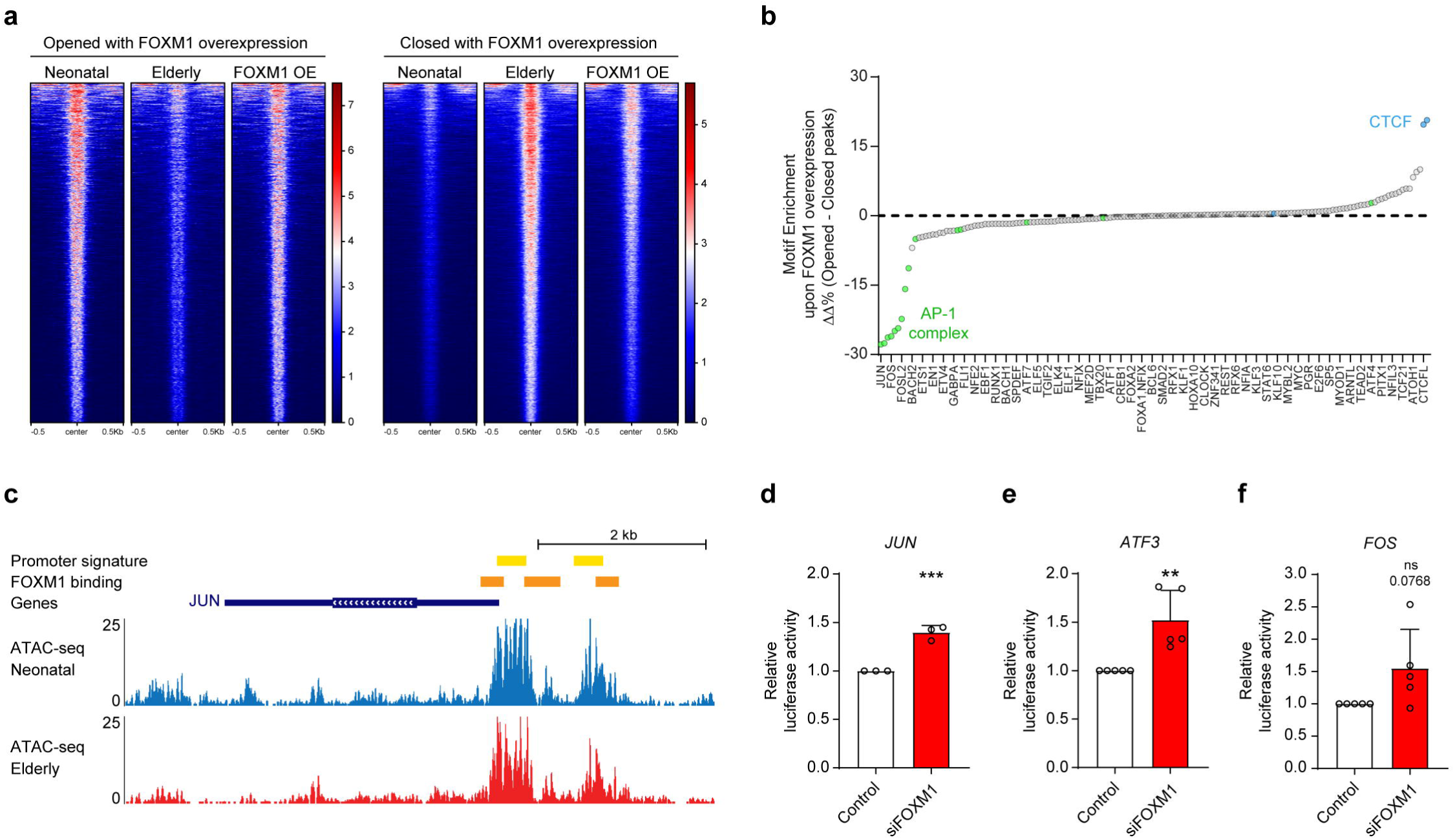
*FOXM1* overexpression rejuvenates the chromatin accessibility profile through repression of AP-1. a.Heatmaps showing the ATAC-seq peak signals rejuvenated by *FOXM1* in neonatal, elderly and *FOXM1*-overexpressing elderly HDFs. Signal is restored from the elderly profile to the neonatal profile by either increasing (chromatin opens, left) or decreasing (chromatin closes, right) upon lentivirus-mediated *FOXM1* overexpression, in elderly HDFs. Each peak was ordered in descending order based on the mean value of ATAC seq signal per region. b.Comparison of HOMER-defined Known Motif Enrichment between opened and closed rejuvenated ATAC-seq reproducible peaks. Positive values represent enrichment in opened peaks (present in neonatal but not in control elderly HDFs), while negative values represent enrichment in closed peaks (not present in neonatal HDFs but present in control elderly HDFs). Members of the AP-1 complex are highlighted in green. Horizontal axis names only every other 3 TFs, for simplicity. c.Genomic tracks representing epigenetic features in the *JUN* locus. Candidate cis regulatory elements with promoter-like signature (Prom sig) from the ENCODE project are shown in yellow. Candidate FOXM1 binding sites at *JUN* promoter (ChIP-seq data from the ENCODE project) are shown in orange. ATAC-seq signal in neonatal (blue) and elderly (red) HDFs showing that chromatin accessibility is kept during aging. d, e, f. Gene expression analysis of *JUN* (b), *ATF3* (c) and *FOS* (d) by RT-qPCR in neonatal HDFs treated with siRNA against *FOXM1*. ** *p*-value ≤0.01; *** *p*-value ≤0.0001 by unpaired Student’s t-test. Values represent mean ± SD.

### FOXM1 is a repressor of the senescence-associated pioneer factor AP-1

We then explored if FOXM1 is a repressor of members of the AP-1 complex. The simpler mechanistic model would be FOXM1 operating via the promoter regions of components of the AP-1 complex. For this reason, we inspected the chromatin accessibility at the promoters of members of the AP-1 complex (Shaulian & Karin, 2002; Garces de Los Fayos Alonso *et al*, 2018). We found the vast majority of these promoters to be accessible in both neonatal and elderly HDFs (Figure 4c; Supplementary Table 8; Supplementary Figure 7), even though the expression of most AP-1 members increases with aging (Supplementary Table 8), concomitantly with reduced *FOXM1* expression.

Thus, we asked whether FOXM1 could control the expression of AP-1 members by acting, directly or indirectly, in their promoters. To this end, we cloned the promoter regions of the *JUN*, *ATF3* and *FOS* genes, encoding AP-1 components previously associated with senescence (Martínez-Zamudio *et al*, 2020; Zhang *et al*, 2021) and found upregulated in elderly cells (Macedo *et al*, 2018), in luciferase reporter constructs. We transfected neonatal HDFs with these constructs and evaluated luciferase expression in response to *FOXM1* knockdown by siRNA (siFOXM1). We found increased luciferase signal in siFOXM1, compared to controls, demonstrating that FOXM1 is indeed a repressor of AP-1 (Figure 4d-f). These results place FOXM1 as an upstream regulator of the AP-1 complex, explaining its ability to control chromatin accessibility during aging and demonstrating that its overexpression reverts age-associated chromatin profiles.

## DISCUSSION

Aging is characterized by subtle alterations of transcriptional profiles, leading to a decline in cellular functions (Sen *et al*, 2016). These transcriptional changes are specific and considered programmed (Stegeman & Weake, 2017). Transcriptional programs are controlled by chromatin accessibility, which changes regulatory landscapes by altering the expression of gene clusters (Buenrostro *et al*, 2015b; Zheng & Xie, 2019). Regulatory landscapes are characterized by many promoter-enhancer interactions allowing for gene cluster transcriptional coordination (Miguel-Escalada *et al*, 2019; Zhu *et al*, 2021). Here we showed that these same mechanistic principles apply to the aging transcriptional programs. We demonstrated that closed and opened chromatin correlate with repression and upregulation of nearby genes’ expression, respectively. These results highlight the potential that TFs and pioneer TFs might have dictating aging transcriptional programs, as previously suggested (Martínez-Zamudio *et al*, 2020; Zhang *et al*, 2021; Di Giorgio *et al*, 2021; Wang *et al*, 2022). We also disclosed that genes differentially expressed during aging tend to be clustered in the genome, exhibiting similar transcriptional changes. We further examined a specific regulatory landscape that contains key genes linked to aging: *FOXM1*, which is gradually repressed along advancing age and whose induction is able to delay aging phenotypes (Macedo *et al*, 2018; Ribeiro *et al*, 2022), and *TEAD4*, a member of the YAP:TEAD pathway whose downregulation drives senescence (Xie *et al*, 2013; Fu *et al*, 2019). Binding motifs for both TFs were enriched in neonatal-specific available chromatin regions. We showed that this regulatory landscape comprises up to 9 genes and is partially conserved in vertebrate genomes, and we demonstrated that at least *TEAD4*, *FOXM1 and RHNO1* are co-regulated by an enhancer that is active in neonatal HDFs and inactive in elderly HDFs. Within this regulatory landscape, other genes (e.g. *FKBP4*, *TULP3* and *TSPAN9*) were similarly downregulated with aging, except *ITFG2*, which was upregulated in elderly HDFs. Interestingly, ITFG2 is a component of the KICSTOR complex which, under catabolic conditions, functions as a negative regulator in the amino acid-sensing branch of mTORC1, known to be activated in aging (Wolfson *et al*, 2017). *FKBP4* encodes for a progestin receptor (PR) co-chaperone repressed with endometriosis and leading to infertility (Hirota *et al*, 2008), and *TULP3* mutations cause multisystem fibrosis originating from disrupted ciliary composition and DNA damage (Devane *et al*, 2022), altogether thus highlighting the functional role of this regulatory landscape in aging processes.

By studying chromatin conformation and accessibility in the genomic landscape of *TEAD4* and *FOXM1*, we found an age-dependent accessible region, named C10-P2, that interacts with the *FOXM1/RHNO1* promoter. We showed this region to be required for proper expression of *FOXM1*, *RHNO1* and *TEAD4*, and its deletion to induce senescence in neonatal HDFs. Although our data indicated that *FOXM1* expression is modulated by multiple accessible loci with enhancer activity in neonatal HDFs, and even though reorganization of topological domains has been associated with *in vitro* senescence (Chandra *et al*, 2015; Zirkel *et al*, 2018; Guan *et al*, 2020), we found the interactome at the *FOXM1/RHNO1* promoter to be largely stable with aging. In contrast, we found many regions with age-dependent accessibility states, in line with previous reports using other cell types and *in vitro* senescence (Moskowitz *et al*, 2017; Ucar *et al*, 2017; Shan *et al*, 2018; Martínez-Zamudio *et al*, 2020; Zhang *et al*, 2021). Notably, while the chromatin accessibility of the *FOXM1/RHNO1* promoter and most *FOXM1* enhancers is maintained in elderly HDFs, the enhancer region C10-P2 becomes inaccessible. Interestingly, the genes co-regulated by the C10 enhancer, *FOXM1*, *RHNO1* and *TEAD4,* have all been associated with cell proliferation (Laoukili *et al*, 2005; Kim *et al*, 2010; Cotta-Ramusino *et al*, 2011; Takeuchi *et al*, 2017; Barger *et al*, 2021), DNA damage response (Cotta-Ramusino *et al*, 2011; Zona *et al*, 2014; Lindsey-Boltz *et al*, 2015; Hazan *et al*, 2019) and senescence (Xie *et al*, 2013; Hernandez-Segura *et al*, 2017; Macedo *et al*, 2018; Fu *et al*, 2019). Our findings demonstrate the pivotal importance of the C10 enhancer in the regulation of a transcriptional hub required for cell homeostasis and survival. The concurrent loss of chromatin accessibility in this enhancer and *FOXM1*, *RHNO1* and *TEAD4* downregulation during aging further suggests that changes in a single, master *cis*-regulatory element contribute to the transcriptional drift towards senescence. The uncovered properties of C10 open new perspectives to ameliorate aging features by modulating the activity of this enhancer using targeted site directed approaches (Han *et al*, 2018; Li *et al*, 2020).

Most strikingly, we found that chromatin regions that are specifically available in neonatal HDFs are enriched for binding motifs of TEAD and FOXM1 TFs, as well as for binding motifs of CTCF TF, which age-related repression leads to defective POLD1 expression and accelerated senescence (Hou *et al*, 2021), and FOXO3 TF, for which genetic polymorphisms have exhibited consistent associations with longevity in diverse human populations (Morris *et al*, 2015). Distinctly, we found that elderly-specific available regions exhibit prevalent binding motifs of the AP-1 complex. These results are concurring with previous studies in induced senescence models, in which AP-1 TFs have been described as pioneer factors that bind and remodel chromatin, thus triggering and maintaining a reversible senescence transcriptional program (Martínez-Zamudio *et al*, 2020; Zhang *et al*, 2021). In this work we show that the expression of genes from the AP-1 complex is repressed by *FOXM1*, a gene highly expressed in neonatal cells and that is transcriptionally downregulated along age. These findings bring novel insight into AP-1 complex regulation during aging, with *FOXM1* repression accounting for the AP-1-driven senescence program. Indeed, previous studies have shown the ectopic expression of *FOXM1* to revert senescence *in vitro* and *in vivo* (Macedo *et al*, 2018; Ribeiro *et al*, 2022). Here we further unveil that ectopic expression of *FOXM1* acts to remodel the chromatin profile of elderly HDFs towards a rejuvenated state. Our findings suggest that these chromatin changes are induced indirectly through the repression of the AP-1 complex. This is in agreement with the reported reversion of senescence transcriptional programs upon AP-1 knockdown (Martínez-Zamudio *et al*, 2020; Zhang *et al*, 2021). Importantly, the promoters of *JUN*, *ATF3* and *FOS*, the three members of the AP-1 complex we examined, are accessible in both neonatal and elderly HDFs, and are thus potentially responsive to FOXM1-mediated repression in an age-independent manner. These observations highlight the potential usage of AP-1 negative regulators, such as *FOXM1*, to revert senescence associated phenotypes in elderly cells, as we have shown by ectopically expressing *FOXM1*. Importantly, these results also suggest that the AP-1 complex is required not only to trigger but also to maintain elderly-specific chromatin profiles and the senescence transcriptional program.

In conclusion, this study advances our understanding of the *FOXM1* key role in aging, strengthening its senotherapeutic potential. We identified a cis-regulatory element whose accessibility is lost during aging and that accounts for *FOXM1* downregulation, as well as for the downregulation of other genes within the regulatory landscape that, altogether, represents a crucial transcriptional hub required to sustain cell fitness and to prevent senescence. In addition to this regulatory landscape, the study further discloses FOXM1 as a repressor of pioneer factors of the AP-1 family, driving (and maintaining) a senescence transcriptional program, and as being able to significantly rescue the changes in chromatin accessibility that account for genome-wide altered regulatory landscapes in aging. Noteworthy, FOXM1 induction does not appear to change indiscriminately the chromatin profile of elderly cells, but rather to mainly revert age specific regions. This grades FOXM1 induction as a safe strategy capable of reprogramming an aging transcriptional program and of reverting aging phenotypes.

## MATERIAL AND METHODS

### Cell culture

Human fibroblasts established from skin biopsies of healthy Caucasian males were acquired from ZenBio (Neonatal: DFM021711A) and NIGMS Human Genetic Cell Repository, Coriell Institute (Neonatal: GM21811, Elderly: AG10884). Only early passage dividing fibroblasts with cumulative population doubling level PDL<24 were used in all experiments. Cells were cultured in Minimal Essential Medium (MEM) with Earle’s salts and L-glutamine (10-010-CV, Corning) supplemented with 15% Fetal Bovine Serum (Gibco) and 1× Antibiotic-Antimycotic (Gibco) in ventilated flasks at 37 °C and humidified atmosphere with 5% CO2.

### Viral Infection

Lentiviruses were produced using the Lenti-X Tet-ON Advanced Inducible Expression System (Clontech), following the protocol described therein. Briefly, HEK293T cells were transfected with packaging plasmids pMD2.G (Addgene #12259) and psPAX2 (Addgene #12260), along with transfer vectors, using Lipofectamine 2000 (Life Technologies, Thermo Scientific, CA, USA). This resulted in the production of lentiviruses carrying either empty pLVX–Tight-Puro (Clontech) or pLVX–Tight-Puro–FoxM1-dNdK (Macedo *et al*, 2018), as well as lentiviruses carrying pLVX–Tet-On Advanced (which expresses rtTA; Clontech). Fibroblasts were co-infected for 5-6 h with empty pLVX–Tight-Puro or FoxM1-dNdK and rtTA-expressing lentiviruses at 2:1 ratio, in the presence of 8 μg/ml polybrene (AL-118, Sigma-Aldrich, MO, USA). After that time, fresh media with 750 ng/ml doxycycline (D9891, Sigma-Aldrich, MO, USA) was added to the cells to induce co transduction. Phenotypes were subsequently analyzed and quantified 68h after doxycycline treatment. Transfection efficiency was monitored by western blotting.

### Western blotting

Lentiviral-infected elderly HDFs were detached with Trypsin, washed thrice with ice-cold PBS and lysed in lysis buffer (150 nM NaCl, 10 nM Tris-HCl pH 7.4, 1 nM EDTA, 1 nM EGTA, 0.5% IGEPAL) with 1x cOmplete, EDTA-free Protease Inhibitor (Roche). Lysates were quantified for protein content by the Lowry Method (DC™ Protein Assay, Bio-Rad, CA, USA). Twenty micrograms of total extract were loaded in SDS-polyacrylamide gels for electrophoresis and transferred onto nitrocellulose membranes for western blot analysis. Membranes were blocked during 1h with TBS (50mM Tris-HCl, 150mM NaCl) containing 5% low-fat milk. Primary antibodies were diluted in TBS containing 2% low fat milk: rabbit anti-FoxM1 (13147, ProteinTech Group, Inc.), 1:1000, and mouse anti-α tubulin (T5168, Sigma-Aldrich, CA, USA), 1:50000. Horseradish peroxidase-conjugated secondary antibodies goat anti-rabbit (SC-2004, Santa Cruz Biotechnology) and goat anti-mouse (SC-2005, Santa Cruz Biotechnology) were diluted at 1:3000 in TBS containing 2% low-fat milk. Detection was done using Clarity Western ECL Substrate reagent (Bio-Rad Laboratories, CA, USA) as per manufacturer’s instructions. Quantitative analysis of protein levels was carried out using a GS-800 calibrated densitometer with ImageLab software (Bio-Rad Laboratories).

### Bisulfite sequencing

DNA samples were collected from cell cultures using the Quick-DNA Miniprep Plus Kit (Zymo Research) and promptly used for bisulfite DNA conversion. The EpiTect Bisulphite Kit (QIAGEN) was used accordingly to the manufacturer’s instructions. The *FOXM1/RHNO1* promoter region was amplified with iMax-II DNA Polymerase (INtRON) and the products cloned into the pCR8/GW/TOPO vector (Invitrogen). Primers used are listed in Supplementary Table 9. Clones were sequenced with Sanger sequencing. Sequencing data was analyzed, and the *p*-value determined with QUMA (Kumaki *et al*, 2008).

### RNA-seq analysis

Previously published RNA-seq datasets derived from the same neonatal and elderly HDFs were used in this study (Macedo *et al*, 2018). Fragment counts (FPM) were normalized to the sample library size and scaled by gene using z-scores, in which 0 corresponds to the mean gene expression across all libraries and the scale represents the gene distance from the mean in standard deviation units. The mean z-score from two neonatal HDF replicates and two elderly HDF replicates is presented in this study.

### ATAC-seq (Assay for Transposase-Accessible Chromatin using sequencing)

ATAC-seq was performed on 50.000 cells according to Buenrostro and colleagues (Buenrostro *et al*, 2015a) with minor modifications. Briefly, cells were detached with Trypsin, washed with ice-cold PBS, lysed in ice-cold fresh lysis buffer (10mM Tris-HCl, pH7.5; 10mM NaCl; 3mM MgCl2; 0.1% IGEPAL) and immediately incubated with homemade Tn5 transposase in TAPS-DMF buffer (Picelli *et al*, 2014) at 37°C for 30 min, followed by proteinase K treatment. Immediately after transposition the samples were purified with MinElute PCR Purification Kit (QIAGEN). We performed qPCR analysis on the samples to determine the appropriate number of PCR cycles to create the library, using 1 µL of tagmented DNA, 0.6 µL of 25 µM Ad Primer 1, 0.6 µL of 25 µM Ad Primer 2 (from 2.1 onwards), used for multiplexing, 1 µL 10x SYBR Green I (ThermoFisher Scientific), 5 µL KAPA HiFi HotStart ReadyMix (KAPA Biosystems) and 1.8 µL H2O HyPure, with the program: 98°C for 45 sec; 72°C for 5 min; 98°C for 30 sec; followed by 30 cycles of 98°C for 10 sec, 63°C for 30 sec, 72°C for 30 sec. The final library was generated using 18 uL of tagmented DNA, 2.5 µL of 25 µM Ad Primer 1, 2.5 µL of 25 µM Ad Primer 2, 25 µL KAPA HiFi HotStart ReadyMix (KAPA Biosystems) and 2 µL H2O HyPure, with the program: 98°C for 45 sec; 72°C for 5 min; 98°C for 30 sec; followed by N cycles of 98°C for 10 sec, 63°C for 30 sec, 72°C for 30 sec; and 72°C for 1 min. N is the number of cycles corresponding to one third of the maximum fluorescence intensity detected in the pilot qPCR. Primers used in this study are listed in Supplementary Table 9. The library was immediately purified using the QIAquick PCR Purification Kit (QIAGEN) and eluted in 20 uL. Successful tagmentation was confirmed with a TAE agarose gel electrophoresis.

Libraries were sequenced on a HiSeq 2500 system, High-Output mode, 2 x 100bp paired-end, at Macrogen, South Korea. Good quality for raw reads of the two replicates (neonatal and elderly conditions) was controlled with FASTQC (Andrews, 2010) (v.0.11.8) and the tagmentation adapters were removed by Skewer (Jiang *et al*, 2014) (v.0.2.1). Reads were mapped to the reference human genome (GRCh38/hg38) using Bowtie2 (Langmead & Salzberg, 2012) (v.2.4.2) with parameters “-X 2000 and --very sensitive”. To avoid clonal artefacts, the duplicated mapped reads were removed using Samtools (Li *et al*, 2009). Mapped reads were filtered by the fragment size (≤120 bp) and mapping quality (≥10) with a custom Python (v.3.7.2) script. To call for enriched regions, MACS2 (Zhang *et al*, 2008) (v.2.2.7) with the parameters “--nomodel, --keep-dup 1, -- llocal 10000, --extsize 74, --shift – 37 and -p 0.07” was used. During all the ATAC-seq analysis, the two replicates were processed independently. Then, we applied the Irreproducible Discovery Rate (IDR) (v.2.0.3) to obtain a confident and reproducible set of peaks based on two replicates (Li *et al*, 2011). Further analysis was performed with the ATAC-seq peaks with a IDR score above 831 (reflecting a *p* value < 0.01). To identify the open chromatin regions, the reproducible peaks were filtered to remove blacklisted regions (Amemiya *et al*, 2019), using BEDTools (v.2.29.2) intersect with default parameters (Quinlan & Hall, 2010). BEDTools intersect and merge were used to identify sample-specific peaks (such as Neonatal specific, Elderly specific or dNdKEN specific) and peaks shared between samples (such as Common between Neonatal and Elderly). Differential accessibility analysis was performed with DESeq2 (v.1.32) (Love *et al*, 2014), with differences considered statistically significant when adjusted *p* value < 0.05 and log2FC > 1. Motif enrichment analysis was performed with HOMER (v.4.11) (Heinz *et al*, 2010), using findMotifsGenome.pl. To determine motif enrichment for each sample the difference between the percentage of sequences with the motif and the percentage of background sequences with the motif was calculated. Different enrichments in different samples were then compared by subtracting the enrichment in one sample by the other and ranking the results by size. Graphs show the gene corresponding to each motif, as defined in HOMER’s motifTable. For visualization of the accessibility profiles, peaks from replicates were first averaged with WiggleTools (v.1.2) (Zerbino *et al*, 2014) mean, and the resulting wig files were converted to bigwig files with WigtoBigWig (v.377) (Kent *et al*, 2010). ATAC-seq heatmaps and profiles were generated with deepTools (v.3.5.1) (Ramírez *et al*, 2016), by first computing a matrix with the parameters “--referencePoint center, -b 500, -a 500, --binSize 10, --sortRegions descend” and the bigwig files abovementioned and by, then, using the tools plotHeatmap and plotProfile, with default parameters. To classify the open chromatin regions identified in this study, BEDTools (v.2.29.2) was used to identify the ATAC-seq peaks whose coordinates have some overlap with candidate cis-regulatory elements (cCREs) derived from ENCODE, using datasets from SCREEN (v.2) (Moore *et al*, 2020).

### Gene clusters and gene expression covariation

To define sets of genes whose expression may be co-regulated by shared CREs, all differently expressed genes between neonatal and elderly HDFs (logFC ≥0.5; *p*-value ≤0.05), as determined by RNA-seq (Macedo *et al*, 2018), were selected. Using grep -Ff, transcription start sites (TSS), from a list of TSS obtained from the Ensembl website (v.105), through the BioMart data mining tool (Kinsella *et al*, 2011; Cunningham *et al*, 2022), were attributed to all differently expressed genes (1436 downregulated and 1873 upregulated). Those genes were grouped with BEDTools cluster, option -d, so that grouping occurred when TSS are no more than 100.000 bp apart. Using Microsoft Excel, the number of genes showing covariation, either down or upregulation, during aging per cluster was assessed, which was then compared to the expected number, assuming no co-regulation, with a Chi-squared test.

### Correlation between chromatin accessibility of CREs and gene expression

To determine the number of putative CREs in the genomic landscapes of genes whose expression is affected by aging, first all putative promoters were excluded from further analyses. BEDTools (v.2.29.2) was used to remove ATAC-seq peaks from our datasets that overlapped with any TSS described in Ensembl (v.105), through the BioMart data mining tool (Kinsella *et al*, 2011; Cunningham *et al*, 2022). Then, grep -Ff was used to attribute genomic coordinates, also obtained with BioMart, to all differently expressed genes during aging (1436 downregulated and 1873 upregulated). Using Microsoft Excel, the gene start coordinate was extended by 10, 20, 30, 50 and 100 kb in each genomic direction. The resulting coordinates constitute what we define as the genomic landscape of genes, based on genetic distance. To determine the number of putative CREs per gene landscape, the list of ATAC-seq peaks without TSS overlap was crossed with the defined genomic landscapes of differently expressed genes during aging using BEDTools (v.2.29.2) intersect, with option -c. The results where then directly processed and plotted in Graphpad (v8.0.2).

### 4C-seq (Circular Chromatin Conformation Capture followed by next-generation sequencing)

4C-seq was performed on about 10 million cells using the enzymes *Dpn*II (first digestion) and *Csp*6I (second digestion), according to van de Werken et al., 2012 (van de Werken *et al*, 2012) with minor modifications. The 4C template was purified using an Amicon Ultra-15 Centrifugal Filter Device (Milipore). At least two libraries were independently prepared with the Expand Long Template polymerase (Roche) using the primers listed in Supplementary Table 9. Libraries were purified with the QIAquick PCR Purification kit (QIAGEN) followed by the Agencourt AMPure XP reagent (Beckman Coulter). Libraries were sequenced on HiSeq 2500 system, Rapid Run mode, 1 x 50bp single-end, at Macrogen. Previously described (Noordermeer *et al*, 2011; Splinter *et al*, 2012) processing was employed with a custom Perl script. More than 1.5 million reads per sample were aligned to the human genome (hg38) using Bowtie (Langmead *et al*, 2009) (requiring unique alignments, -m 1). Reads within fragments flanked by restriction sites of the same enzyme (checked with BEDTools) (Quinlan & Hall, 2010) or fragments smaller than 40bp were filtered out. Mapped reads were then converted to reads-per-first-enzyme fragment end units and smoothed using a 30 fragment mean running window algorithm. For visualization, the smoothed reads from each sample were normalized to 1 million reads per kilobase (https://github.com/porchard/normalize_bedgraph).

### Luciferase assays

Putative enhancer sequences and putative promoter sequences were defined based on our ATAC-seq signal and ChIP-seq data (TF clusters and H3K27ac) from the ENCODE project. Sequences were amplified with iMax-II DNA Polymerase (INtRON) from genomic DNA of neonatal HDFs (DFM021711A). Primers used are listed in Supplementary Table 10. Putative enhancer sequences were TA cloned into pCR8/GW/TOPO vector (Invitrogen) and subcloned into pGL4.23-GW (Pasquali *et al*, 2014) (Addgene #60323) using Gateway LR Clonase II (Gateway cloning, Invitrogen). The empty pGL4.23-GW was used as negative control. The TK promoter from pGL4.54[luc2_TK] (Promega #E5061) was cut with *Kpn*I and *Hind*III, ligated into the *Kpn*I/*Hind*III-digested pME-MCS (Kwan *et al*, 2007) (Tol2 kit #237) and subcloned into pGL4.23-GW as above. pNL1.1PGK[Nluc/PGK] (Promega #N1441) was used as transfection control. Cells were transfected in a 6-well plate with 1 pmol of pGL4.23-GW and 0.61 fmol of pNL1.1PGK[Nluc/PGK] per well with Lipofectamine 3000 (Invitrogen) according to manufacturer’s instruction. Cells were seeded 12-24h before transfection. Luciferase activity was assessed 48h post-transfection using the Nano-Glo Dual-Luciferase Reporter Assay System (Promega) in a Synergy 2 Multi-Mode Microplate Reader (BioTek). The enhancer activity is expressed as the luc2/Nluc ratio (pGL4.23 GW/pNL1.1PGK[Nluc/PGK]), and is presented as a relative response ratio, in which the activity of the empty vector and positive control vector were normalized to 0 and 1, respectively. The enhancer activity of putative enhancers was compared to the control with a one-way analysis of variance (ANOVA) with Dunnett’s correction for multiple comparisons (** *p*-value ≤0.01, **** *p*-value ≤0.0001), in GraphPad Prism 8.

To test the response of the *JUN*, *ATF3* and *FOS* gene promoter sequences to siRNA against *FOXM1*, the *PGK* promoter from pNL1.1.PGK[Nluc/PGK] (Promega #N1441) was removed by digestion with *Kpn*I and *Hind*III, and the purified plasmid backbone was used for Gibson cloning. The pGL4.54[luc2_TK] (Promega #E5061) was used as transfection control. pNL1.1.PGK[Nluc/PGK] was used as positive control. A multiple cloning site was inserted in the *Kpn*I/*Hind*III-digested pNL1.1.PGK[Nluc/PGK], recreating the empty vector pNL1.1[Nluc], which was used as negative control. Putative promoter sequences were selected based on the Eukaryotic Promoter Database EPD (Dreos *et al*, 2017) and our ATAC-seq data. Sequences were amplified with iMax-II DNA Polymerase (INtRON) from genomic DNA of neonatal HDFs (DFM021711A) and Gibson cloned into the *Kpn*I/*Hind*III-digested pNL1.1.PGK[Nluc/PGK]. Primers used are listed in Supplementary Table 11. Cells were seeded in a 24-well plate 12-24h before transfection. Cells were transfected with 50nM of a siRNA against *FOXM1* (SASI_Hs01_00243977, Sigma-Aldrich) or a negative control siRNA (siRNA Universal Negative Control #1, Sigma-Aldrich), using Lipofectamine RNAiMax (Invitrogen), following manufacturer’s instructions. About 24h later, cells were transfected with 0.5 pmol of pGL4.54[luc2_TK] and 0.05 pmol of the promoter-test vectors per well with Lipofectamine 3000 (Invitrogen). Luciferase activity was assessed 48h post-transfection (72h post siRNA transfection) using the Nano-Glo Dual-Luciferase Reporter Assay System (Promega) in a Synergy 2 Multi-Mode Microplate Reader (BioTek). The promoter activity is expressed as a normalized Nluc/luc2 ratio (pNL1.1.AP1[Nluc/AP1]/pGL4.54[luc2_TK]), in which the ratio of each promoter in cells treated with the control siRNA was normalized to 1, in each independent experiment. The conditions were compared with an unpaired Student’s *t*-test using GraphPad Prism 8 (* *p*-value ≤0.05, ** *p*-value ≤0.01, *** *p*-value ≤0.001, **** *p*-value ≤0.0001). Error bars represent mean ± s.d.

### CRISPR/Cas9-mediated deletions and FACS-sorting

sgRNA spacer sequences upstream and downstream of the target regions were selected on Benchling (Benchling [Biology Software], 2018) based on metrics from Doench and colleagues(Doench *et al*, 2016) and Hsu and colleagues (Hsu *et al*, 2013). The oligonucleotide sequences can be found in Supplementary Table 12. Pairs of oligonucleotides were ordered (Sigma-Aldrich) and annealed, followed by cloning in the *BbsI*-digested pSpCas9(BB)-2A-GFP (PX458) (Addgene, #48138) and the pU6 (BbsI)_CBh-Cas9-T2A-mCherry (Addgene, #64324) vectors. Neonatal HDFs were seeded in 6-well plates and transfected with 500ng of each plasmid per well using Lipofectamine 3000 (Invitrogen) according to manufacturer’s instructions. After 72h, cells were sorted in a FACSAria II cell sorter (BD Biosciences) using an 85μm nozzle and the blue (488nm) and yellow/green (561nm) lasers. Cells were gated by forward scatter area (FSC-A) vs. side scatter area (SSA-A) and FSC-A vs. FSC-height (FCS-H) plots to exclude dead cells and doublets/clumps, respectively. The gates were established based on the autofluorescence of mock controls. 500 GFP and mCherry double-positive cells were collected into 96-well plates to establish polyclonal cell cultures. Non-transfected (mock) cells and cells transfected with the empty vectors were always sorted, collected, and grown in parallel to the CRISPR/Cas9-targeted cells. When reaching 75%-95% cell confluency, cells were consecutively transferred to a 48-well, 12-well, 6-well plate and ultimately to a T25 flask, from which cells were used for DNA extraction and genotyping, RNA extraction and RT-qPCR analysis, immunostaining assays, SA-β-gal activity assay and live-cell imaging.

### Genotyping

CRISPR/Cas9-mediated genomic deletions were validated in the cultures used for the phenotypic analyses described below. Resuspended cells were washed with 10mM Tris HCl pH 8, incubated in Tris buffer at 99°C for 15 min, and treated with 1µg/µL proteinase K at 56°C for 30 min, followed by proteinase K inactivation at 95°C for 10 min. The extracted DNA was used for genotyping with iMax-II DNA Polymerase (INtRON) following manufacturer’s instructions and adjusting the extension time for each locus. Mock cells and cells transfected with the empty vectors were genotyped in parallel. Primers used are listed in Supplementary Table 13. Efficient deletion of the different loci was determined by the presence of an amplicon with the expected size with a TAE agarose gel.

### Real-time quantitative PCR

Total RNA was extracted using TRIzol reagent (Invitrogen), treated with DNAse I (Thermo Scientific), and precipitated with sodium acetate in ethanol. iScript Synthesis Kit (Bio-Rad) was used for cDNA synthesis. RT-qPCR was performed with iTaq Universal SYBR Green Supermix (Bio-Rad) in a CFX96 or a CFX384 Touch Real-Time PCR Detection System (Bio-Rad) according to manufacturer’s instructions. Primers used are listed in Supplementary Table 14. Data were analyzed using the CFX Maestro software (Bio-Rad). Non-reverse transcribed and blank controls were included. Three technical replicates were used per target gene. Expression was normalized to the *TBP* and *HPRT1* housekeeping genes, and different experimental samples were normalized to the mean expression of the control samples. The samples were compared with an unpaired Student’s *t*-test using GraphPad Prism 8 (* *p*-value ≤0.05, ** *p*-value ≤0.01, *** *p*-value ≤0.001, **** *p*-value ≤0.0001).

### Immunostaining

Cells were seeded 2-3 days before fixation in 8-well µ-slides (Ibidi) or 96-well CellCarrier Ultra microplates (PerkinElmer). When at 75-95% confluency, were fixed in 4% paraformaldehyde (Electron Microscopy Sciences) in PBS for 15 min, permeabilized with 0.3% Triton-X100 in PBS for 7 min, washed 3 times in 0.05% Tween 20 in PBS (PBS-T) for 5 min, blocked in 10% fetal bovine serum (FBS) in PBS-T for 1h at RT and incubated with primary antibodies in 5% FBS in PBS-T, overnight at 4°C. The primary antibodies used in this study were anti-Ki67 (8D5, #9449, Cell Signaling, at 1:500), anti-53BP1 (#4937, Cell Signaling, at 1:500), anti-p21 (F-5, sc-6246, Santa Cruz, 1:800) and anti p16INK4a (ab7962, Abcam, at 1:200). After washing 3 times in 0.05% Tween 20 in PBS (PBS-T) for 5 min, cells were incubated with secondary antibodies in 5% FBS in PBS-T, at 1:1500, for 45 min at RT. The secondary antibodies used in this study were anti-rabbit Alexa 488 and anti-mouse Alexa 568 (Life Technologies). Samples were then washed 5 times in PBS-T for 5 min, counterstained with DAPI (Sigma-Aldrich, at 1:1000) for 10 min at RT and washed in PBS for 5 min. Images were acquired in a Leica DMI6000 B (Leica Microsystems) equipped with a Hamamatsu FLASH4.0 (Hamamatsu) camera. Images were then analyzed using Fiji – ImageJ (Schindelin *et al*, 2012). The samples were compared by unpaired Student’s *t*-test using GraphPad Prism 8 (* *p*-value ≤0.05, *** *p* value ≤0.001).

### SA-β-gal activity assay

Cells were seeded 3 days before the assay. Cells were incubated with 1nM Bafilomycin A1 (Sigma-Aldrich) for 90 min, to induce lysosomal alkalinization. The fluorogenic substrate for β-galactosidase, DDAOG (Setareh Biotech), was then added to the cell culture to a final concentration of 10µM and cells were incubated for 90 min. Cells were then sorted in a FACSAria II cell sorter (BD Biosciences) using an 85μm nozzle and the red (633nm) laser. All cells within an experiment were detected using the same voltage settings. Cells were gated by forward scatter area (FSC-A) vs. side scatter area (SSA-A) and FSC-A vs. FSC-height (FCS-H) plots to exclude dead cells and doublets/clumps, respectively. The gates were established based on the autofluorescence of non-treated cells. Analysis of FACS data was done using FlowJo v10 software (FlowJo, LLC). The samples were compared by unpaired Student’s *t*-test using GraphPad Prism 8 (** *p*-value ≤0.01).

### Time-lapse live-cell imaging

Cells were seeded 24h before image acquisition in 4-well or 8-well µ-slides (Ibidi). Cells were kept under controlled temperature, CO2 and humidity levels during the experiments and were at 50-75% confluency when imaged. Image acquisition took 6-12h and each field was imaged every 2.5-4min. Time-lapse images were either acquired in a Leica DMI6000 B (Leica Microsystems) or in a Nikon TI (Nikon) inverted microscope, equipped with the cameras Hamamatsu FLASH4.0 (Hamamatsu) or Andor iXon 888 (Andor), respectively. Images were then analyzed using Fiji – ImageJ (Schindelin *et al*, 2012). Mitotic duration was determined as the time (min) from nuclear envelop breakdown (NEB) to anaphase onset (AO). The populations were compared by unpaired Student’s *t*-test using GraphPad Prism 8 (**** *p*-value ≤0.0001).

### Statistical analysis

Details about statistical analysis and software used can be found in figure legends and respective method description. Values represent mean ± s.d. Statistical analysis was performed with GraphPad Prism 8.

## DATA AVAILABILITY STATEMENT

The 4C-seq data sets for this publication were deposited in the European Nucleotide Archive (ENA) at EMBL-EBI under the accession number PRJEB46917. ATAC-seq data will be publicly available in a later stage. Bioinformatic datasets and any additional information required to reanalyze the data reported in this paper is available from the lead contacts upon request.

## SUPPLEMENTARY DATA

**Supplementary Figure 1. Multiple comparisons of chromatin accessibility variation during aging.**

a. Comparison of HOMER-defined Known Motif Enrichment between a subset of neonatal and elderly-specific open chromatin regions that presented significant quantitative differences of ATAC-seq signal (darker blue and red), determined by DESeq2. Negative values represent bigger enrichment in neonatal peaks, while positive values represent bigger enrichment in elderly peaks. Horizontal axis names only every other 9 TFs, for simplicity.

b. Comparison of HOMER-defined Known Motif Enrichment between all age-specific and age-enriched, determined by DESeq2, neonatal and elderly ATAC-seq peaks. Negative values represent bigger enrichment in neonatal peaks, while positive values represent bigger enrichment in elderly peaks. Horizontal axis names only every other 9 TFs, for simplicity.

c. Quantification of ATAC-seq reproducible peaks whose accessibility does not change between neonatal and elderly cells, in genes that are up (red) or downregulated (blue) during aging. Values represent mean ± sd. *p*-value was determined by Mann-Whitney U test.

**Supplementary Figure 2. The regulatory landscape of *TEAD4*, *FOXM1* and *RHNO1*.**

a. Heatmap of expression levels of genes in the putative regulatory landscape containing *TEAD4*, *FOXM1* and *RHNO1*, interrogated from RNA-seq datasets of neonatal and elderly HDFs (Macedo *et al*, 2018). Z-score column color intensities represent higher (red) to lower (blue) mean gene expression in two replicates of each age. Expression of *NRIP2* and *TEX52* was not detected.

b. Bisulfite sequencing of the *FOXM1/RHNO1* promoter in neonatal and elderly HDFs. Each circle represents a CpG. Black-filled circles represent methylated CpGs.

c. 4C-seq analysis of the *FOXM1/RHNO1* promoter interactions with active promoters in its proximal genomic landscape. Purple bars match the purple-colored genes shown in a, representing promoters interacting with the *FOXM1/RHNO1* promoter.

**Supplementary Figure 3. Phylogenic conservation of the *FOXM1* and *RHNO1* locus.**

a. Graphical representation of the order and orientation of homologous genes around *FOXM1* in different ancestral reconstructions of Chordata genomes, adapted from the PhyloView tool of the Genomicus Vertebrates (v109.01) database, using data from Ensembl. Each color represents a different gene. An asterisk (*) is used to indicate *RHNO1*-like genes that have been described in non-Euteleostomi species, although they are not annotated as such in Genomicus.

b. Genomic tracks from the UCSC browser representing transcript annotations and epigenetic features in the *BL11002* locus (red), a predicted *FOXM1* homologous gene, and the nearby gene *BL00819* (blue) in the lancelet *Branchiostoma lanceolatum*. Annotations, transcriptomics and epigenetic data from Marlétaz et al. (Marlétaz *et al*, 2018). Transcriptomics and epigenetic data tracks correspond to 8 hpf *B. lanceolatum* embryos. First track represents Aligned Ensembl Human proteins. Only two representative *B. lanceolatum* transcripts for each gene are shown. H3K27ac and ATAC-seq signals suggest an active regulatory region between the transcription starting sites of both genes. RNA-seq data suggest both genes are actively expressed in 8 hpf embryos. The locus was identified with specific primers designed to amplify *FOXM*, a *FOXM1* homologous gene in the lancelet *Branchiostoma floridae,* which perfectly amplify the *BL11002* gene. Gene *BL00819* presents an inferred ontology with the terms “DNA damage checkpoint signaling” and “cellular response to ionizing radiation” in the Ensembl Metazoa database (Yates *et al*, 2022), terms consistent with the described role of *RHNO1* in humans (Cotta-Ramusino *et al*, 2011). UCSC Genome Browser on European amphioxus (v. Bl71nemh 20/11/13) (BraLan2), region Sc0000191:2,367-15,655.

c. Genomic tracks from the UCSC browser representing gene annotations and genomic data in the locus of *LOC109482876* and *LOC109482936* in *Branchiostoma belcheri*. Gnomon, an NCBI eukaryotic gene prediction tool (Souvorov *et al*, 2010), predicts *LOC109482876* as a *FOXM1*-like gene and *LOC109482936* as an *RHNO1*-like gene. No clear CpG island is identified between *LOC109482876* and *LOC109482936*. UCSC Genome Browser on Haploidv18h27 Apr. 2016 Belcher’s lancelet (GCF_001625305.1), region NW_017804254.1:1,230,170-1,244,604.

d. Genomic tracks from the UCSC browser representing gene annotations in the locus of *foxm* and *E3 ubiquitin-protein ligase CCNB1IP1-like* in the vase tunicate *Ciona intestinalis*. ENSCING00000019370 is the sole *FOXM* gene annotated in this species by the NCBI Reference Sequence (RefSeq) collection (Pruitt *et al*, 2005) and Ensembl (Cunningham *et al*, 2022), and it is identified as a *FOXM1* homologous gene in Genomicus Vertebrates (v109.01) (Nguyen *et al*, 2022). Although ENSCING00000009823, identified as a *E3 ubiquitin-protein ligase CCNB1IP1-like* by UniProt (Bateman *et al*, 2023), is not described as a *RHNO1*-like gene, RHNO1 is known to bind RAD18, an E3 ubiquitin ligase, in human cells (Cotta-Ramusino *et al*, 2011). UCSC Genome Browser on C. intestinalis Apr. 2011 (Kyoto KH/ci3), region chr4:5,001,156-5,011,537.

e. Phylogenetic tree representing the evolutionary relationships between different taxonomic groups, with example species, and the genome structure encompassing the locus of *FOXM1* homologous genes, based on data from Genomicus Vertebrates (v109.01), Amphioxous (v01.01) and Tunicates (v03.01) (Nguyen *et al*, 2022) and taxonomic classifications by UniProt (Bateman *et al*, 2023). We identified the *FOXM1* homologue in *B. lanceolatum* using primers previously designed for *B. floridae*’s *FOXM* gene (Yu *et al*, 2008). Genomicus Amphioxous (v01.01) (Nguyen *et al*, 2022) identified the human *RHNO1* as a homologue gene for *BL00819* in *B. lanceolatum*. Gnomon (Souvorov *et al*, 2010) predicts *LOC109482876* as a *FOXM1*-like gene and *LOC109482936* as an *RHNO1*-like gene. *C. intestinalis*’s ENSCING00000019370 is annotated as *FOXM* by the NCBI Reference Sequence (RefSeq) collection (Pruitt *et al*, 2005) and Ensembl (Cunningham *et al*, 2022). The NCBI Reference Sequence (RefSeq) collection (Pruitt *et al*, 2005) annotates *FOXM1* and *RHNO1* genes in *Callorhinchus milii*, *Erpetoichthys calabaricus* and *Homo Sapiens,* although *Callorhinchus milii*’s *RHNO1* homologue is not annotated in Ensembl/Genomicus. An asterisk (*) is used to indicate *RHNO1*-like genes that have been described in non-Euteleostomi species.

**Supplementary Figure 4. Age-specific analysis of chromatin conformation and accessibility in the genomic landscape of *TEAD4*, *FOXM1* and *RHNO1*.**

a. 4C-seq analysis of chromatin interactions with the *FOXM1/RHNO1* promoter in neonatal (dark blue) and elderly (purple) HDFs.

b. ATAC-seq analysis of chromatin accessibility in the genomic landscape of *FOXM1/RHNO1* in neonatal (light blue) and elderly (red) HDFs. Replicates were used to perform peak calling and identify age-specific and common accessible genomic regions (black bars). Orange bars indicate putative enhancer regions examined in this study.

c-e. Chromatin accessibility in neonatal and elderly HDFs in enhancer C2, c, enhancer C5, d, and enhancer C9, e.

**Supplementary Figure 5. Deletion of enhancer C10 leads to decreased Ki67 positive cells, increased 53BP1/p21-positive cells and mitotic delay.**

a,b. Percentage of cells staining positive for the Ki67 proliferation marker in mock, Cas9 only treated and C10-P2-deleted cells.

c,d. Percentage cells with double-positive staining for 53BP1 DNA damage and p21/CDKN1A cell cycle arrest markers in mock, Cas9-only treated and C10-P2-deleted cells.

e. Mitotic duration (nuclear envelop breakdown, NEB, to anaphase onset, AO) in mock, Cas9-only treated and C10-P2-deleted cells.

* p-value ≤0.05, ** p-value ≤0.01, *** p-value ≤0.001, **** p-value ≤0.0001 by unpaired Student’s t-test. All values represent mean ± SD. Scale bar: 20 µm.

**Supplementary Figure 6. Validation of *FOXM1* overexpression and its impact on chromatin accessibility.**

a,b. FOXM1 protein levels in neonatal HDFs infected with lentivirus carrying empty pLVX–Tight-Puro vector or pLVX–Tight-Puro–FoxM1-dNdK. FOXM1 levels were normalized to α-tubulin levels. Only the levels of endogenous full-length FOXM1 were quantified (note that FOXM1 binds its own promoter, therefore ectopic FOXM1-dNdKEN induces endogenous FOXM1).

c. Venn diagram showing overlap of the ATAC-seq reproducible peaks from elderly and FOXM1-dNdKEN-elderly HDFs. FOXM1-dNdKEN expression induces loss of accessibility in 25.016 peaks and gain of accessibility in 14.899 new regions.

d. Venn diagram showing overlaps of the ATAC-seq reproducible peaks from neonatal, elderly and FOXM1-dNdKEN-elderly HDFs. Upon FoxM1-dNdKEN ectopic expression, 17.752 of the previously identified elderly-specific accessible genomic regions (red outline) were closed (red), and 6.488 genomic regions previously identified in the neonatal-specific dataset (blue outline) were opened (turquoise).

**Supplementary Figure 7. Members of the AP-1 complex retain chromatin accessibility during aging.**

Genomic tracks representing epigenetic features in the locus of 10 members of the AP-1 complex, representing the 4 subfamilies (JUN, FOS, ATF, MAF). Candidate cis regulatory elements with promoter-like signature (Prom sig) from the ENCODE project are indicated by yellow bars. Candidate sites of FOXM1 and TEAD4 binding inferred from ENCODE ChIP-seq data are indicated by orange bars. ATAC-seq signal in neonatal (blue peaks) and elderly (red peaks) HDFs showing chromatin accessibility of the genomic regions nearby the promoters of members of the AP-1 complex.

**Supplementary Table 1.** Intersection of ATAC-seq peaks with candidate cis-regulatory elements from ENCODE.

**Supplementary Table 2.** HOMER Known Motif Enrichment in all neonatal and elderly peaks, and comparison.

**Supplementary Table 3.** HOMER Known Motif Enrichment in neonatal-specific and elderly-specific peaks, and comparison.

**Supplementary Table 4.** HOMER Known Motif Enrichment in neonatal-specific, elderly specific peaks, and common but DESeq2 enriched peaks, and comparison.

**Supplementary Table 5.** Gene clusters determined by distance between TSS.

**Supplementary Table 6.** Genomic coordinates and features of the putative cis regulatory elements selected in this work and studied in luciferase assays.

**Supplementary Table 7.** HOMER Known Motif Enrichment in neonatal-specific and elderly-specific peaks, opened or closed upon FOXM1 overexpression, respectively, and comparison.

**Supplementary Table 8.** Chromatin accessibility and RNA-seq data related to members of the AP-1 complex during aging.

**Supplementary Table 9.** Primers used for bisulfite sequencing (BS-seq), 4C-seq and ATAC-seq.

**Supplementary Table 10.** Primers used to amplify the putative enhancer regions.

**Supplementary Table 11.** Primers used to amplify the promoter sequences of members of the AP-1 complex, for luciferase reporter assays.

**Supplementary Table 12.** sgRNA spacer sequences with overhangs for cloning used for CRISPR/Cas9-mediated deletions.

**Supplementary Table 13.** Primers used to genotype cells edited with CRISPR/Cas9.

**Supplementary Table 14.** Primers used for RT-qPCR.

## ACKNOWLEDGMENTS

We are thankful to Juan Tena (CABD, CSIC-UPO, Seville) for his kindness sharing custom Perl scripts for 4C-seq data processing. The authors acknowledge the support of the i3S Scientific Platforms: BioSciences Screening; Translational Cytometry; Cell Culture and Genotyping; and Advanced Light Microscopy, member of PPBI (POCI-01-0145-FEDER-022122).

## Author Contributions

FJF performed all wet lab experiments and respective analyses. FJF and MG performed computational analyses and data interpretation. JT performed preliminary ATAC-seq plotting and computational analysis. JB and EL supervised the project and contributed to all analyses. FJF, JB and EL wrote the manuscript with input from all authors. All authors contributed to the development and discussion of the work. All authors have read and agreed to the final version of the manuscript.

