## Supplementary figures and images for "FOXM1 expression reverts aging chromatin profiles through repression of the senescence-associated pioneer factor AP-1"

### Supplementary Figure 1

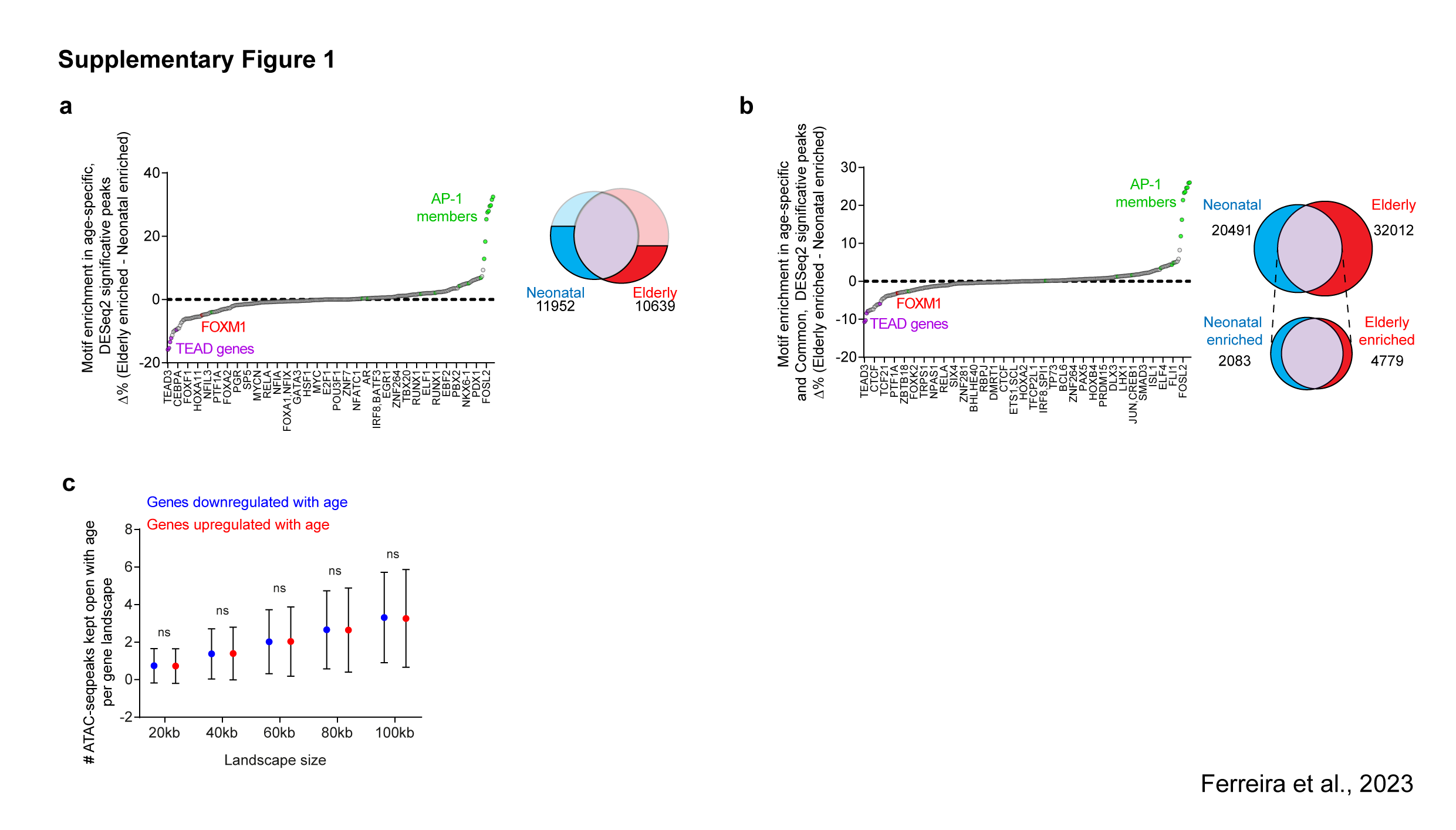

### Supplementary Figure 2

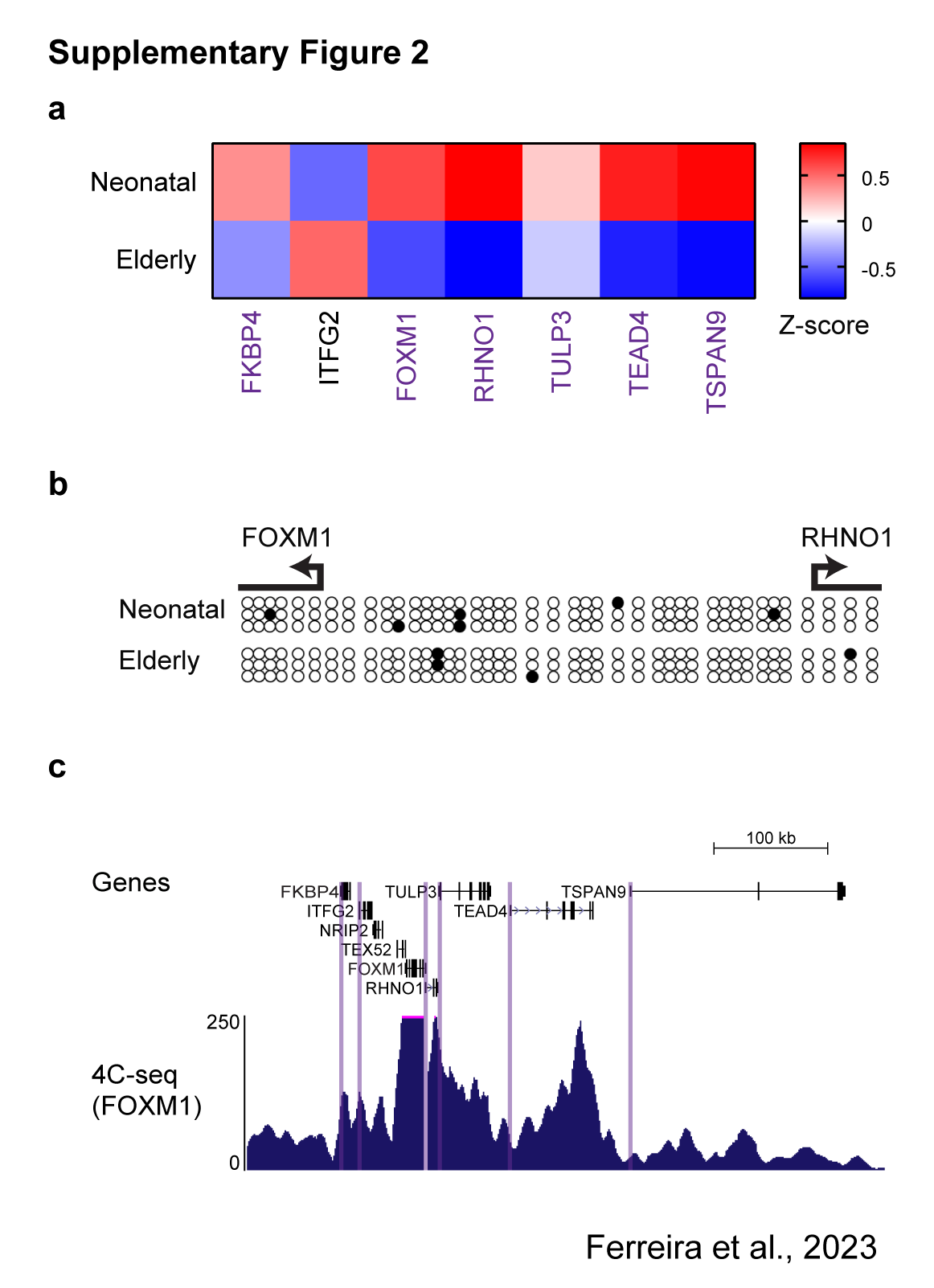

### Supplementary Figure 3

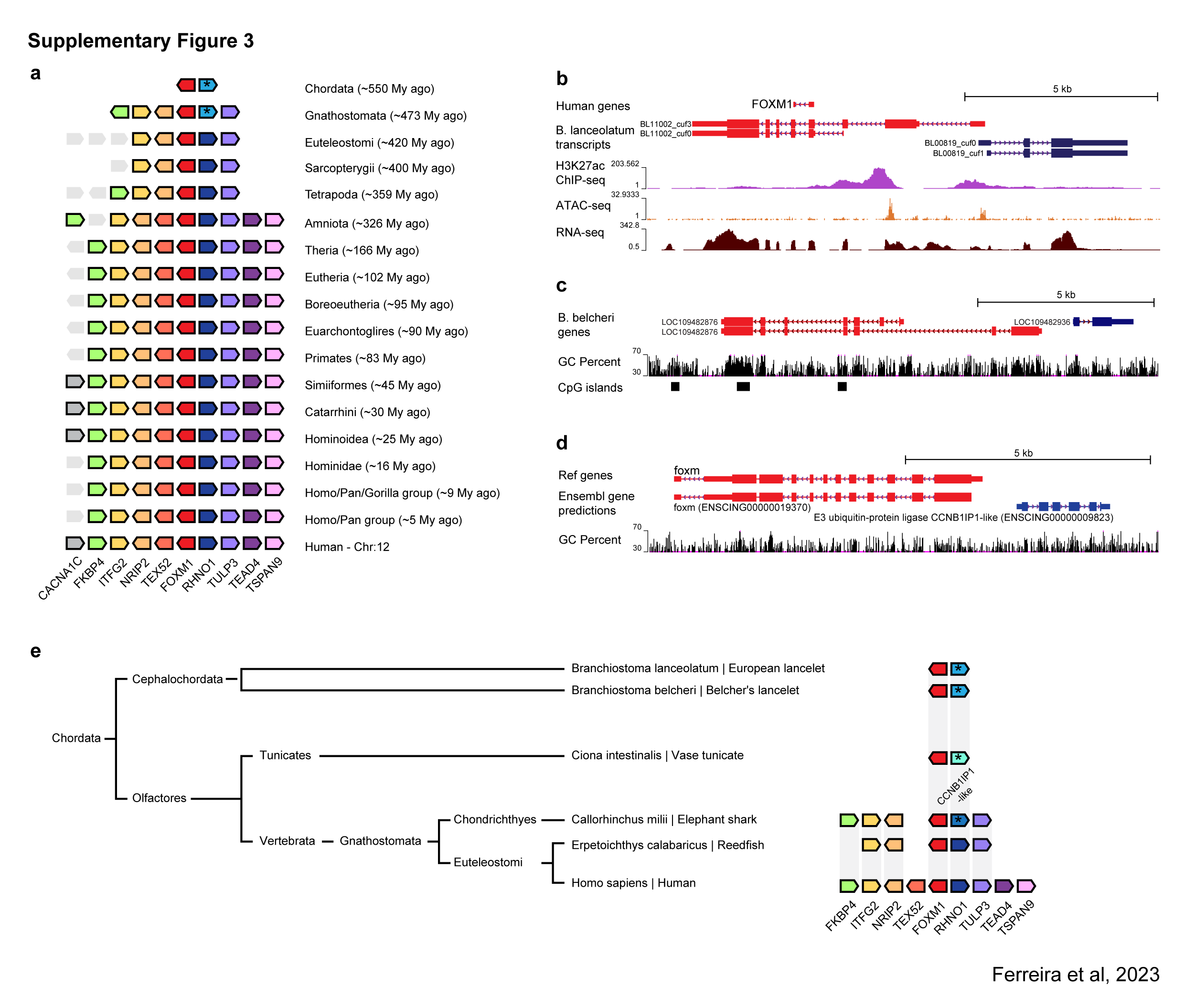

### Supplementary Figure 4

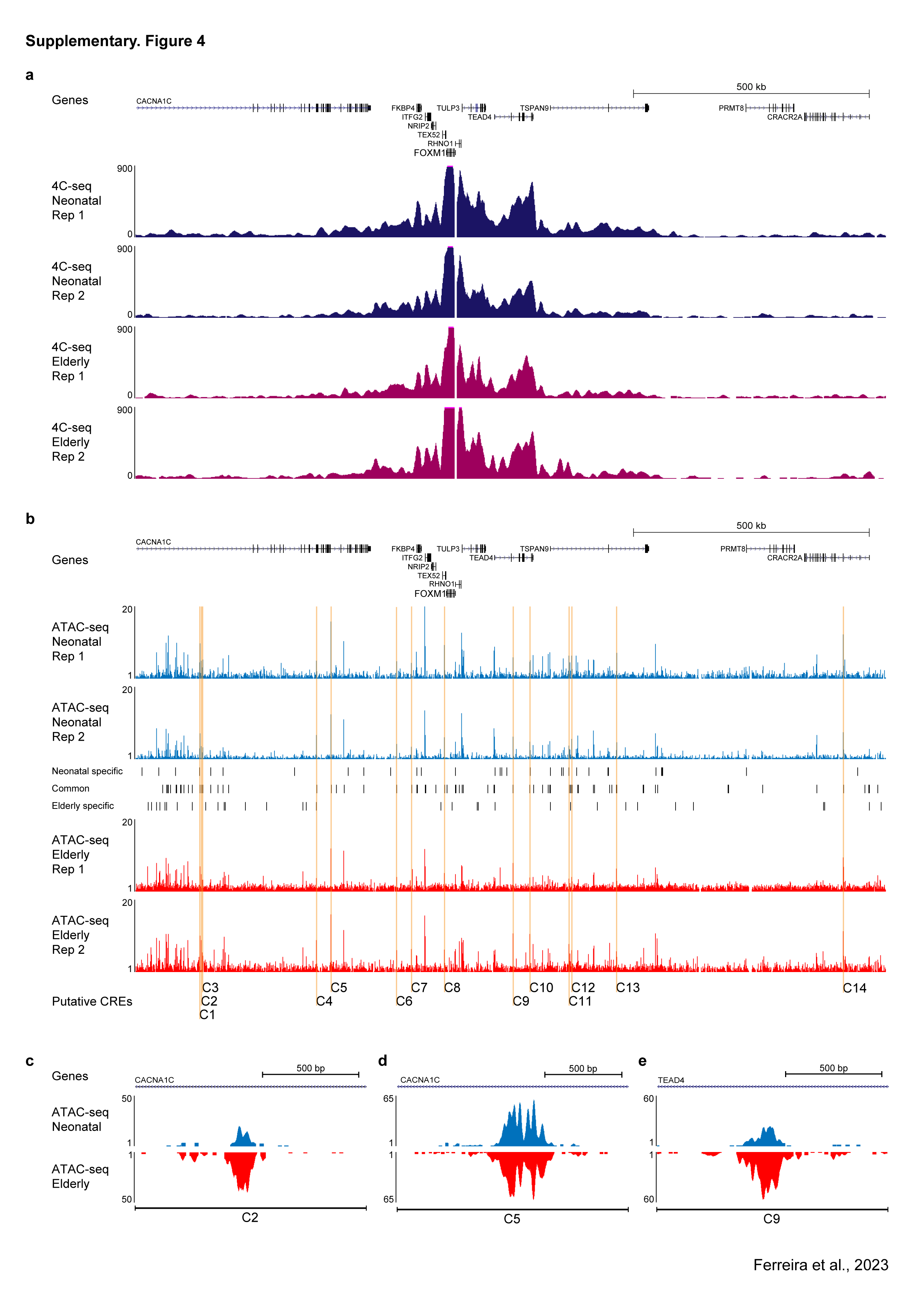

### Supplementary Figure 5

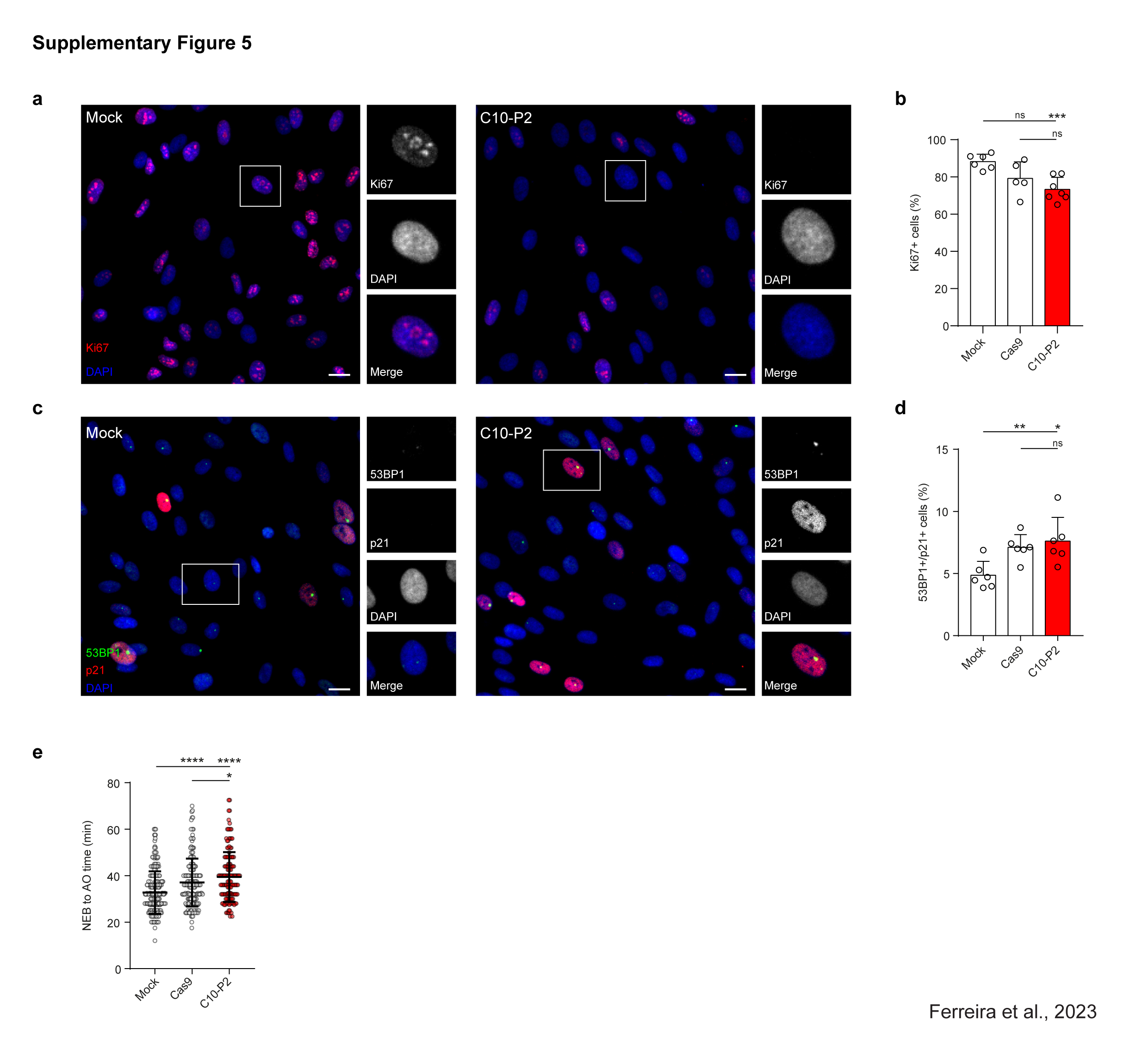

### Supplementary Figure 6

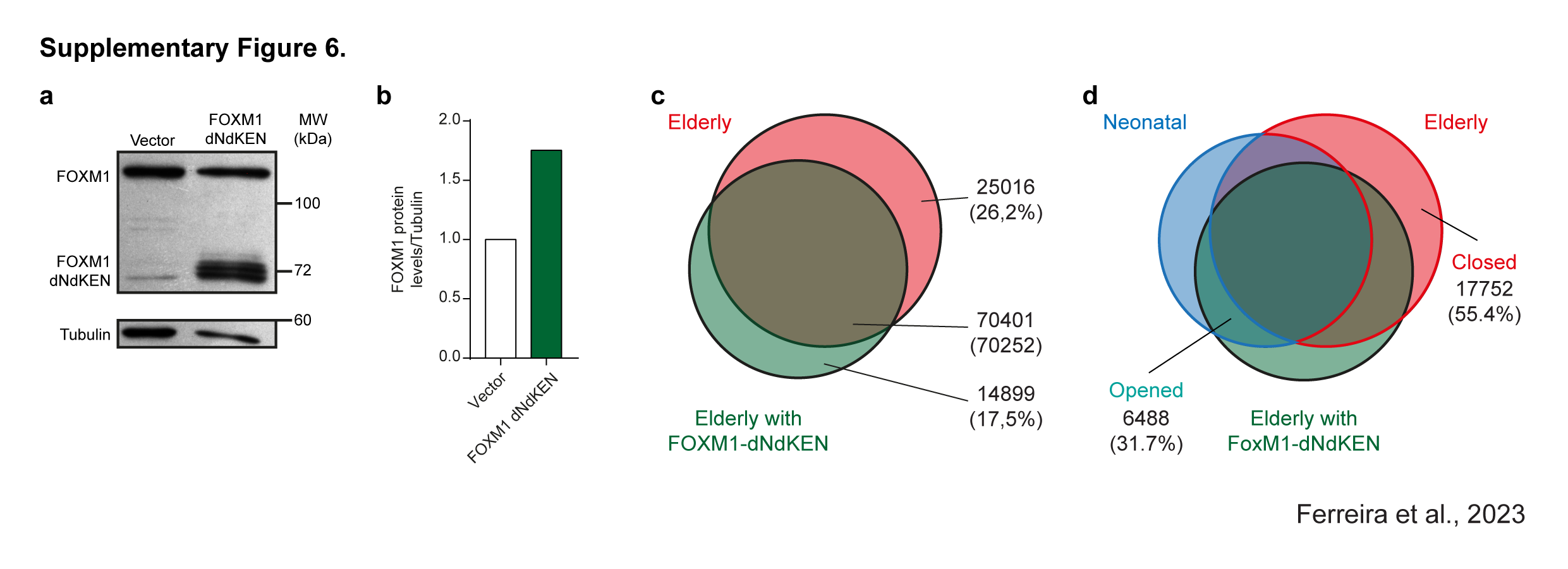

### Supplementary Figure 7

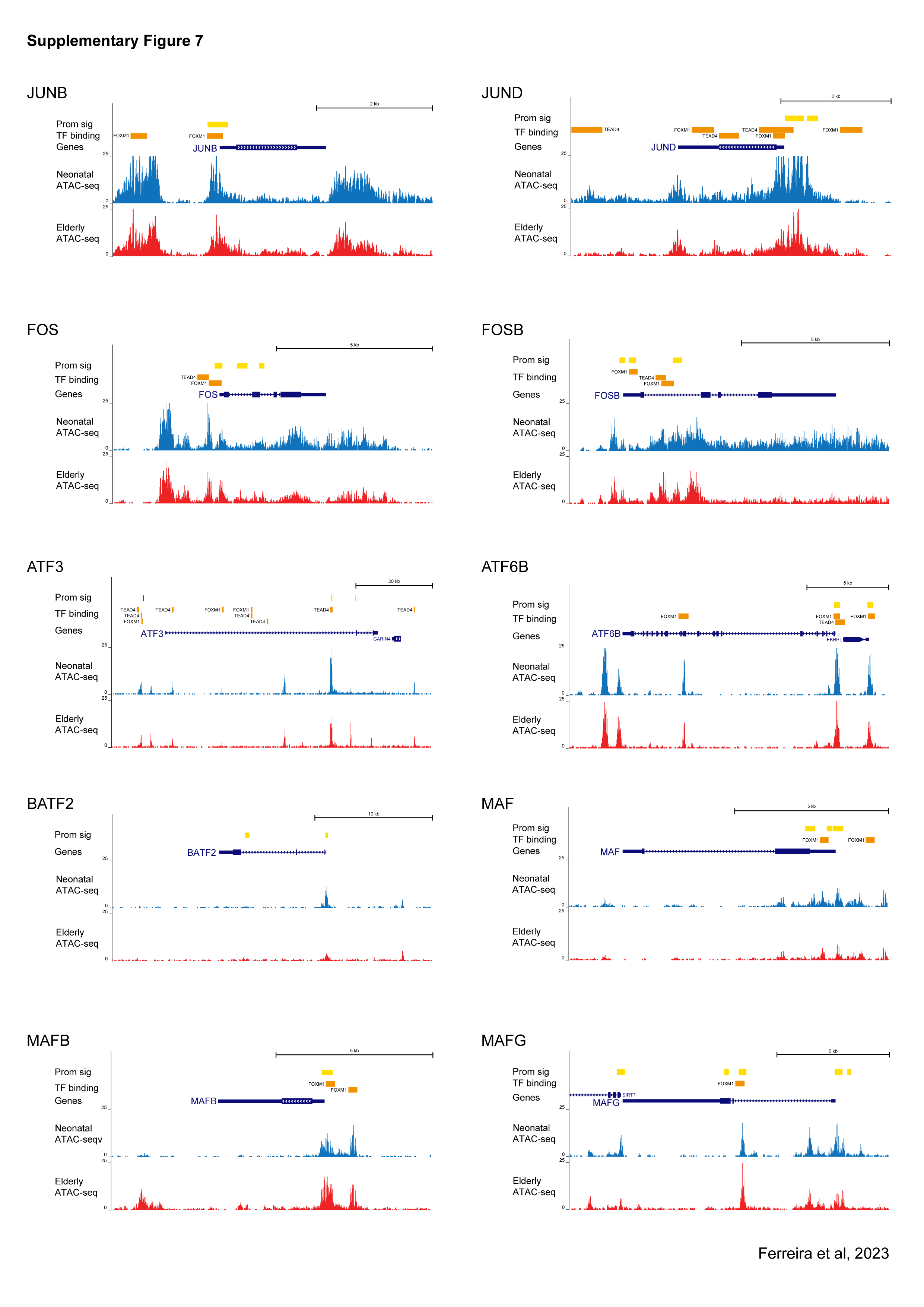
