## Supplementary Tables 9 to 14 for "FOXM1 expression reverts aging chromatin profiles through repression of the senescence-associated pioneer factor AP-1"

**Supplementary Table 9.** Primers used for bisulfite sequencing (BS-seq), 4C-seq and ATAC-seq.

| **Assay** | **Primer** | **Sequence (5´-3´)** |
|---|---|---|
| BS-seq | FOXM1/RHNO1_fw | TATTTCGTTTATTTTTAGATTGTAG |
| | FOXM1/RHNO1_rv | AAACCGCAACTCCTAACAAACC |
| 4C-seq | LNonread_ foxm1 | CAAGCAGAAGACGGCATACGAggctttagttgatttcctcactggg |
| | LRead_foxm1_BC0 | AATGATACGGCGACCACCGAACACTCTTTCCCTACACGACGCTCTTCCGATCTccctcccctcccgggatc |
| | LRead_foxm1_BC1 | AATGATACGGCGACCACCGAACACTCTTTCCCTACACGACGCTCTTCCGATCTGTGccctcccctcccgggatc |
| | LRead_foxm1_BC2 | AATGATACGGCGACCACCGAACACTCTTTCCCTACACGACGCTCTTCCGATCTAAGccctcccctcccgggatc |
| | LRead_foxm1_BC3 | AATGATACGGCGACCACCGAACACTCTTTCCCTACACGACGCTCTTCCGATCTTCAccctcccctcccgggatc |
| ATAC-seq | Ad1 | AATGATACGGCGACCACCGAGATCTACACTCGTCGGCAGCGTCAGATGTG |
| | Ad2.1 | CAAGCAGAAGACGGCATACGAGATTCGCCTTAGTCTCGTGGGCTCGGAGATGT |
| | Ad2.2 | CAAGCAGAAGACGGCATACGAGATCTAGTACGGTCTCGTGGGCTCGGAGATGT |
| | Ad2.4 | CAAGCAGAAGACGGCATACGAGATGCTCAGGAGTCTCGTGGGCTCGGAGATGT |

**Supplementary Table 10.** Primers used to amplify the putative enhancer regions enhancers.

| Putative Enhancer | Primer | Sequence (5´-3´) |
|---|---|---|
| C1 | E2336123_fw | CAAAGCATGGATGGAGAAGC |
| | E2337234_rv | GCCTCTCAGGGACTACTGACC |
| C2 | E2338768_fw | GTGGAATAGGGCTCTTGAGG |
| | E2340128_rv | AAAACAGCTTCGTGCTCACC |
| C3 | E2341758_fw | CATTCAGGGGTAACTGACTGG |
| | E2342421_rv | GAGGAAGAGGATGAATGAATGC |
| C4 | E2582858_fw | GATGGGATCATGGCTTATGG |
| | E2583813_rv | GGAAGAGGGGCTTTGTCC |
| C5 | E2613113_fw | CCAGCTCGGCCTTTGAATCC |
| | E2614507_rv | GCCTGAAAATTTTGGAGCTGC |
| C6 | E2752309_fw | TCCCCAGTCGCTTAAAACCC |
| | E2753433_rv | CGTTATCCTGCTTCCCTCCC |
| C7 | E2783481_fw | TGTAGCATCACACTCAGCCC |
| | E2784969_rv | TGGTTCAGCAAAAGCCAGG |
| C8 | E2853209_fw | CAGGGAAGCAGGGAGAAGC |
| | E2854479_rv | ACAGTTCCAACCCCATCACC |
| C9 | E2998545_fw | GCCTGGAGGAAGAGTGAGG |
| | E2999781_rv | GCGAGAAGTTGTTCGAATCC |
| C10 | E3033857_fw | GGTGGACAGGAAATGTCAGG |
| | E3035537_rv | TCTCCTATTTTCTCGAAGTCACC |
| C11 | E3116634_fw | ATTTGCTTGGCTGTTTCTGC |
| | E3118210_rv | GCTGTTAGTGCTGGAAATGAGG |
| C12 | E3122669_fw | CGCATCTGTCTGAATGTGC |
| | E3123934_rv | CTGCACAGCTCCTTTTAAACC |
| C13 | E3218162_fw | GCCCTGGTTTTCTCTAGGC |
| | E3219021_rv | GACACAACAGAAAACAGACATCC |
| C14 | E3697913_fw | GCAATGTGATGTGGAAGAGG |
| | E3699199_rv | ACAACTGGGTCTCAGGAAGG |

**Supplementary Table 11.** Primers used to amplify the promoter sequences of members of the AP-1 complex, for luciferase reporter assays.

| Gene | Primer | Sequence (5´-3´) |
|---|---|---|
| *JUN* | JUN_prom_fw | <u>TGGCCTAACTGGCCGGTACC</u>GGAGTTCCGTGATTTGCTTGC |
| | JUN_prom_rv | <u>AGTACCGGATTGCCAAGCTT</u>TCTCTGGACACTCCCGAAAC |
| *ATF3* | ATF3_prom_fw | <u>TGGCCTAACTGGCCGGTACC</u>CTTCCCAGCCTCACCTAGTC |
| | ATF3_prom_rv | <u>AGTACCGGATTGCCAAGCTT</u>GGCGGGCTGTTTGTCTGG |
| *FOS* | FOS_prom_fw | <u>TGGCCTAACTGGCCGGTACC</u>GTTCTCTCTCATTCTGCGCC |
| | FOS_prom_rv | <u>AGTACCGGATTGCCAAGCTT</u>AGAGCTGGGTAGGAGCACG |

Underlined sequences correspond to overhangs inserted for subsequent Gibson cloning into the *Kpn*I/*Hind*III-digested pNL1.1.PGK[Nluc/PGK], equivalent to the pNL1.1[Nluc] backbone.

**Supplementary Table 12.** sgRNA spacer sequences with overhangs for cloning used for CRISPR/Cas9-mediated deletions.

| Putative Enhancer | sgRNA | Forward sequence (5´-3´) | Reverse sequence (5´-3´) |
|---|---|---|---|
| C2 | Upstream1 | CACCGATGGGTATGCTAGCTCAACT | AAACAGTTGAGCTAGCATACCCATC |
| | Upstream2 | CACCGGCTCTTGAGGGCTTATCCT | AAACAGGATAAGCCCTCAAGAGCC |
| | Downstream1 | CACCGCAGATGAATTATGAACACC | AAACGGTGTTCATAATTCATCTGC |
| | Downstream2 | CACCGTATAATGTGGAATCCTGTGA | AAACTCACAGGATTCCACATTATAC |
| C5 | Upstream1 | CACCGTACCCGCAGAACACTTGACA | AAACTGTCAAGTGTTCTGCGGGTAC |
| | Upstream2 | CACCGCTCAAACAATTAAGTGCCC | AAACGGGCACTTAATTGTTTGAGC |
| | Downstream1 | CACCGACAGTTTAGACCACTCGCAA | AAACTTGCGAGTGGTCTAAACTGTC |
| | Downstream2 | CACCGTTTACACCATAGATGTACCC | AAACGGGTACATCTATGGTGTAAAC |
| C8 | Upstream1 | CACCGAGAAGCCTGGGTCCCCACAA | AAACTTGTGGGGACCCAGGCTTCTC |
| | Upstream2 | CACCGAGTGATCAGTCTCCTTTGTG | AAACCACAAAGGAGACTGATCACTC |
| | Downstream1 | CACCGCACAGTGAGGTCGGTACCTG | AAACCAGGTACCGACCTCACTGTGC |
| | Downstream2 | CACCGTCAGCTCCCTCACCGATGAA | AAACTTCATCGGTGAGGGAGCTGAC |
| C9 | Upstream1 | CACCGTCCTTGAAGGGACCCTTGAA | AAACTTCAAGGGTCCCTTCAAGGAC |
| | Upstream2 | CACCGCAGAGGAACGCCAAGCTCGG | AAACCCGAGCTTGGCGTTCCTCTGC |
| | Downstream1 | CACCGAAATGAAAGCGGCCGTGAGC | AAACGCTCACGGCCGCTTTCATTTC |
| | Downstream2 | CACCGTCGGCCTAGCCTAGCGTGC | AAACGCACGCTAGGCTAGGCCGAC |
| C10 | Upstream1 | CACCGCAGGCACTGAGGACGAGTCG | AAACGCAGGCACTGAGGACGAGTCG |
| | Upstream2 | CACCGAACACACCAGCCGGCGACG | AAACCGTCGCCGGCTGGTGTGTTC |
| | Downstream1 | CACCGAAACAGGAGCCCGAAAGCCT | AAACAGGCTTTCGGGCTCCTGTTTC |
| | Downstream2 | CACCGTGCCGGACTAGATTTAACTG | AAACCAGTTAAATCTAGTCCGGCAC |
| | Middle1 | CACCGTGTGTCTGGAAGAGCTGGG | AAACCCCAGCTCTTCCAGACACAC |
| | Middle2 | CACCGAAGAATCCAGTGTGCCAAGC | AAACGCTTGGCACACTGGATTCTTC |

**Supplementary Table 13.** Primers used to genotype cells edited with CRISPR/Cas9.

| Enhancer | Primer | Sequence (5´-3´) |
|---|---|---|
| C2 | E2338667_fw2 | TAACTTTGGAGCAGCGAGCG |
| | E2340304_rv2 | AGGCTTTCTTAAGTGCCTTTTGGC |
| C5 | E2613113_fw* | CCAGCTCGGCCTTTGAATCC |
| | E2614507_rv* | GCCTGAAAATTTTGGAGCTGC |
| C8 | E2853161_fw3 | CCAGGGTCAGCACCTATCAG |
| | E2854494_rv3 | ACTGGAGTGAAAGCTGTAGCC |
| C9 | E2998431_fw2 | GAAGCAGACTGAAAATGGTTTGG |
| | E2999945_rv3 | AGGAAGACTCGGGAAAGTCAAG |
| C10, C10-P1, C10-P2 | E3033815_fw2 | GTAGCTTGTTTTGGGTGGTTGC |
| | E3036030_rv2 | AGTCTCCCCACTCTGTCTCC |

*same shown in Supplementary Table 9.

**Supplementary Table 14.** Primers used for RT-qPCR.

| Gene | Primer | Sequence (5´-3´) |
|---|---|---|
| *FOXM1* | FOXM1_fw | ATACGTGGATTGAGGACCACT |
| | FOXM1_rv | TCCAATGTCAAGTAGCGGTTG |
| *RHNO1* | RHNO1_fw | GAAGGCTGTGCGGTCTGC |
| | RHNO1_rv | CAATTCCAACCCATGCCAGC |
| *TEAD4* | TEAD4_fw | CAGGAAGCAGGTCTCCAGC |
| | TEAD4_rv | TTAGCTTGGCCTGGATCTCG |
| *TBP* | TBP_fw | GAGCCAAGAGTGAAGAACAGTC |
| | TBP_rv | GCTCCCCACCATATTCTGAATCT |
| *HPRT1* | HPRT1_fw | CCTGGCGTCGTGATTAGTGAT |
| | HPRT1_rv | AGACGTTCAGTCCTGTCCATAA |
